# Population history and genetic adaptation of the Fulani nomads: Inferences from genome-wide data and the lactase persistence trait

**DOI:** 10.1101/650986

**Authors:** Mário Vicente, Edita Priehodová, Issa Diallo, Eliska Podgorná, Estella S. Poloni, Viktor Černý, Carina M. Schlebusch

## Abstract

Human population history in the Holocene was profoundly impacted by changes in lifestyle following the invention and adoption of food-production practices. These changes triggered significant increases in population sizes and expansions over large distances. Here we investigate the population history of the Fulani, a pastoral population extending throughout the African Sahel/Savannah belt. Based on genome-wide analyses we propose that ancestors of the Fulani population experienced admixture between a West African group and a group carrying both European and North African ancestries. This admixture was likely coupled with newly adopted herding practices, as it resulted in signatures of genetic adaptation in contemporary Fulani genomes, including the control element of the *LCT* gene enabling carriers to digest lactose throughout their lives. The lactase persistence (LP) trait in the Fulani is conferred by the presence of the allele T-13910, which is also present at high frequencies in Europe. We establish that the T-13910 LP allele in Fulani individuals analysed in this study lies on a European haplotype background thus excluding parallel convergent evolution. Our findings further suggest that Eurasian admixture and the European LP allele was introduced into the Fulani through contact with a North African population/s. We furthermore confirm the link between the lactose digestion phenotype in the Fulani to the *MCM6/LCT* locus by reporting the first Genome Wide Association study (GWAS) of the lactase persistence trait. We also further explored signals of recent adaptation in the Fulani and identified additional candidates for selection to adapt to herding life-styles.

## Introduction

The Fulani are a large and widely dispersed group of both nomadic herders and sedentary farmers living in the African Sahel/Savannah belt. Currently, they reside mostly in the western part of Africa, but some groups are dispersed up to the Blue Nile area of Sudan in the east (Stenning 1957; Delmet 2000). Although some historians postulated an origin of the Fulani in ancient Egypt or the Upper Nile valley (Lam 2001), written records suggest that the Fulani spread from West Africa (currently Senegal, Guinea, Mauritania) around 1,000 years ago, reaching the Lake Chad Basin 500 years later (Newman 1995; Fay 1997). They founded several theocratic states such as Massina (Ba and Daget 1962), Sokoto (Johnston 1967), or Takrur (McIntosh et al. 2016), and many Fulani abandoned the nomadic lifeway and settled down, including in large urban centers. This expansion was accompanied by a process of group absorption of sedentary peoples called *Fulanisation*, that led to shifts in ethnic identity of some sedentary peoples, as has been described in North Cameroon (Schultz 1984). However, several Fulani groups retained the very mobile lifestyle relying on the transhumance of their livestock and cattle milking. These fully nomadic or at least semi-nomadic groups are still present in several Sahelian locations, especially in Mali (de Bruijn and van Dijk 1995), Niger (Dupire 1962), Central African Republic (Boutrais 1988) and Burkina Faso (Riesman 1974; Benoit 1982). All Fulani speak the *fulfulde* Niger-Congo west-Atlantic language (a language continuum of various dialects), consistent with their postulated Western African ancestry (Greenberg 1963).

Similarly to other pastoralists, the Fulani experienced specific selection pressures probably associated with a lifestyle characterized by transhumance and herding (Podgorna et al. 2015; Triska et al. 2015). Lactase Persistence (LP) is a widely studied genetic trait with evidence of recent selection in populations who adopted pastoralism and heavily rely on dairy products, especially drinking fresh milk (Tishkoff et al. 2007; Schlebusch et al. 2012a; Breton et al. 2014; Macholdt et al. 2014; Liebert et al. 2017). LP is associated with the control element of the *LCT* gene on chromosome 2 (Durham 1992; Hollox et al. 2001; Swallow 2003; Bersaglieri et al. 2004; Mulcare et al. 2004;

Tishkoff et al. 2007; Itan et al. 2010; Ranciaro et al. 2014). Specific polymorphisms in this region prevent the down-regulation of the *LCT* gene during adulthood and confer the ability to digest lactose after weaning (Tishkoff et al. 2007; Ranciaro et al. 2014; Liebert et al. 2017). The LP trait is particularly frequent in northern European populations, pastoralists from East Africa, farmers and pastoralists from the Arabian Peninsula, and Arab speaking pastoralists from northeastern Africa and the Sahel/Savannah belt (Priehodova et al. 2014; Liebert et al. 2017; Priehodova et al. 2017; Schlebusch and Jakobsson 2018; Schlebusch 2019). To date, five different variants conferring LP in populations across the globe have been identified. The independent genetic backgrounds of these polymorphisms suggest convergent adaptation in populations with dairy-producing domesticated animals.

The T-13910 allele is reported to be the key variant regulating maintenance of *LCT* gene expression in European adults. This variant is generally not detected in most East African and Middle Eastern populations, where other LP variants are observed instead (Enattah et al. 2008; Ranciaro et al. 2014; Priehodova et al. 2017; Schlebusch and Jakobsson 2018; Schlebusch 2019). Fulani populations living mainly in the western Sahel/Savannah belt, however, carry the European-LP mutation with frequencies ranging from 18% to 60% (Lokki et al. 2011; Ranciaro et al. 2014; Haber et al. 2016; Cerny et al. 2018). The presence of this “European” LP variant at relatively high frequencies across different Fulani populations is puzzling and could either result from convergent evolution in both Africa and Europe or from gene flow between ancestors of the Fulani and Europeans. The later hypothesis is supported by the fact that T-13910 has not been detected (or is only present at very low frequencies) in neighbouring populations of the Fulani (Ranciaro et al. 2014; Cerny et al. 2018) and that European admixture in Fulani genomes has been reported in previous studies (Henn et al. 2012; Triska et al. 2015).

Details surrounding the European admixture event and the post-admixture selection of the European LP mutation in Fulani genomes remain unclear. Studies based on uniparental markers reported higher frequencies of western Eurasian and/or North African mitochondrial DNA (mtDNA) and Y chromosome haplogroups in the Fulani than in neighbouring populations (Cerny et al. 2006; Cerezo et al. 2011; Buckova et al. 2013; Kulichova et al. 2017). However, studies on *Alu* insertions did not lead to similar results, connecting instead the Fulani with East African pastoralists (Cizkova et al. 2017).

In this study we analyse genome-wide SNP data from 53 Fulani pastoralists from Ziniaré, Burkina Faso to investigate the history of the Fulani population and the patterns of Eurasian admixture in their genomes, and to uncover the origin of the LP variant they carry. We perform genome-wide selection scans to investigate the strength of selection on the LP region and to identify other additional genomic regions that experienced selection during processes of adaptation to herding lifestyles in the Fulani. Lastly, we attempt to identify additional genomic regions associated with the ability of digesting milk during adult life by performing, for the first time in published research, a genome-wide association study (GWAS) on the lactose tolerance phenotype in adults.

## Subjects and Methods

### Sampling

Sampling in Burkina Faso was conducted in collaboration with the Burkinabe CNRST (*Centre national de la recherche scientifique et technologique*) institution in Ouagadougou. Measurements of LP phenotypes and collection of saliva (Oragene kit, DNA Genotek) of the Fulani (FZR, n = 56) were carried out in three nomadic camps located, at the time, northeast of Ziniaré (longitude –1.241395; latitude 12.620579) with research permit No 0495 and help of local assistants. Informed consent was obtained from all the participants included in the study before samples were collected. Additional DNA and LP phenotype measures were collected for a comparative dataset of 63 unrelated volunteers based in Prague with Czech or Slovak (CS) nationality. All Czech and Slovak volunteers signed informed consent on anonymous use of their sample. The study was approved by the Ethical Committee of the Charles University in Prague (approval no. 2016/07) and by the Swedish Ethical Review Authority (approval no. 2019-00479).

### Phenotype test

For estimation of lactase activity, we used the lactose tolerance test (LTT), which is based on the measurement of an increase of blood glucose (glycemia) after consumption of 50 g of lactose on an empty stomach (Arola 1994; Tishkoff et al. 2007). Blood glucose was measured by eBsensor (Visgeneer Inc.). Volunteers were asked to starve overnight (minimally 8 hours) and their base-line blood glucose was measured afterwards. Then they were asked to drink 50 g of lactose dissolved in 200 ml of water (which is equivalent to the amount of lactose in 1 to 2 litres of cow’s milk) (Arola 1994; Tishkoff et al. 2007). Blood glucose was measured 20, 40 and 60 minutes after ingestion. The maximal difference from the base-line from these three measurements was used in genotype-phenotype comparisons.

### Sequencing of 359-bp fragment in intron 13 of the *MCM6* gene

To detect which LP mutations were present in the 56 Fulani and 63 Czech/Slovak genomes, we Sanger sequenced intron 13 of the *MCM6* gene with a previously reported set of primers (Coelho et al. 2009). The primers cover a 359-bp long fragment where 5 known LP associated variants are located. The PCR products were Sanger sequenced by Macrogen (Korea).

### Genome-wide SNP typing

A subset of 55 Fulani and 7 Czech/Slovak individuals were selected for genome-wide genotyping on the Illumina Omni2.5-Octo BeadChip (which contains the T–13910 SNP). The data was aligned to the Human Genome built version 37.

Data management and quality filtering was carried out using PLINK v.1.90 software (Chang et al. 2015). A total of 2,608,742 SNPs were obtained from the 62 individuals. All individuals passed 0.15 data missingness threshold. We subsequently filtered to keep only autosomal SNPs with a SNP missingness filter of 0.1. To account for possible genotyping errors, we applied a Hardy-Weinberg equilibrium filter (HWE) that excluded 90 SNPs (for p ≤ 1e-4). AT and CG SNPs were excluded to prevent strand flipping errors when merging with comparative datasets. Relatedness was measured by identical by state (IBS) analysis and two Fulani individuals were excluded due to potential genetic relatedness. A total of 2,359,821 SNPs and 60 individuals were kept for the study.

The newly generated data will be made available for academic research use through the ArrayExpress database accession number XXXX.

### Merging with previously published data

We merged the new data with published comparative datasets following the same quality control criteria as described above. We added 1295 samples from 39 populations (full descriptive list in Table S1).

For the first dataset (dataset A) we compiled selected groups from the 1000 genomes project (Auton et al. 2015) and a Sahelian dataset (Triska et al. 2015) to merge with the newly generated data. After merging and quality filtering, dataset A had 785 individuals and 1,968,522 autosomal SNPs. This dataset was used in initial analyses of the genetic affinities, selection scans, GWAS and the local ancestry analyses using RFMix involving two reference population sources in the Fulani. We added additional samples to dataset A in order to get a more representative picture of past demographic history by compiling dataset B, which covers 1355 individuals from 41 populations and 297,954 autosomal SNPs (Table S1). Dataset B was used in studies of genetic affinities (such as PCA, admixture analyses and f-statistic methods) and local ancestry analyses using three reference population groups. Dataset B was extended with additional

Eurasian populations for f3-statistics with the Fulani European-like segments (see European-specific analysis and F-statistics based methods section). Furthermore, a San population from Namibia (Schlebusch et al. 2012b) was also included as outgroup in the demographic models using qpGraph (see European-specific analysis and F-statistics based methods section). Datasets were phased using SHAPEIT software (O’Connell et al. 2014), using the HapMap II recombination map.

### Population structure analyses

Population structure analyses were performed on dataset B. We generated a Principal Component Analysis (PCA) that compares Fulani individuals to comparative groups of dataset B. The PCA analyses were performed with EIGENSOFT (Patterson et al. 2006; Price et al. 2006) under default settings. We inferred admixture fractions with ADMIXTURE (Alexander et al. 2009) to investigate relationships among individuals. The number of clusters (K) was set from 2 to 10, replicated 20 times. The cluster-inference and visual inspection was made with Pong v.1.4.5 (Behr et al. 2016).

### Estimating admixture dates

To estimate the time of possible admixture events we used a linkage disequilibrium (LD) decay based method. The date estimations were done for dataset B using Malder, ADMIXTOOLS package (Patterson et al. 2012). The HapMap II genetic map was used as recombination map.

### Local ancestry analyses

We use the RFMix software (Maples et al. 2013) to identify local ancestries of genomic fragments in Fulani genomes. An initial RFMix run was performed with two ancestral populations, represented by 50 YRI, for West African ancestry, and 50 CEU individuals, to represent European ancestry on a total 1,968,522 autosomal SNPs. We ran RFMix analyses with two extra iterations to account for admixture in the source populations and to minimize assignment errors. We set 5 minimum reference haplotypes per tree node and the number of generations to admixture to 30. We used HapMap II genetic map as recombination map and positions outside the map windows were excluded. We ran an additional RFMix analysis with similar settings and added 30 Mozabite individuals as a third parental source to account for potential North African ancestry in Fulani genomes using a total of 297,954 autosomal SNPs.

### Haplotype plots and networks

We extracted the haplotypes of approximately 1.1 Mb (positions 135,759,095 to 136,824,836) surrounding the –13910*T (position 136,608,646) on chromosome 2 from our phased dataset. The selected region was chosen based on RFMix results and the haplotypes were sorted by position of –13910*T/A. We used the same selected region to construct a median joining network using the NETWORK software package ver. 5.0.0.1 employing the median joining (MJ) algorithm (Bandelt et al. 1999) and the maximum parsimony (MP) option (Polzin and Daneschmand 2003). The parameter “Frequency > 1” was activated, so that unique haplotypes are not shown in network plots.

### European-specific analysis and F-statistics based methods

To test the affinity of the European-specific fragments in the Fulani genomes we firstly kept only the European-like regions of the Fulani genomes based on the output of RFMix for dataset B. *f3*-outgroup and pair-wise *fst* were obtained using ADMIXTOOLS. To avoid affinity bias towards CEU (European parental population in RFMix), the individuals previously selected for representing the European source in RFMix were replaced by the remaining 50 individuals of the CEU population in the 1000 genomes project.

To inspect the frequency of a European-like (CEU) fragment being flanked by North-African-like (Mozabite) fragment we recorded the number of European fragments flanked by a North-African ancestry region in relation to the total number of European fragments across all Fulani datasets. To test whether North African regions flank European regions more often than expected, we performed a bootstrap test by randomly selecting fragments across the genome (equal to the number of European-like fragments) and test the likelihood of a random fragment being flanked by a North African ancestry region. This analysis was repeated 9,999 times.

We also tested if the different ancestry distributions across the genome could be explained by two independent admixture events. We simulated 53 phased individuals containing a conservative number of 10,000 haplotypes. These individuals hold similar admixture fractions to the Fulani (0.13 European, 0.19 North African and 0.68 West African). We tested if the three different ancestries follow a random distribution across the genomes, as expected under a neutrality model assumption. This simulation test was performed 100 times and the frequency of a European-like fragment being surrounded by North African-like fragments was recorded.

Lastly we tested population models with specific admixture events using qpGRAPH package (Patterson et al. 2012). The model predictions were performed using Fulani, CEU, Mozabite and Yoruba individuals, using default parameters.

### GWAS

The Fulani (FZR) and Czechs & Slovakian (CS) individuals were used in a genome-wide association scan (GWAS) analysis between their LTT phenotype and their genomic composition (samples extracted from dataset A). The analysis was performed using the GenABEL v. 1.7-6 R package (Aulchenko et al. 2007). A SNP cut-off rate of 0.95, minimum allele frequency of 0.1, a cut-off p-value for the HWE of 1e-08 and false discovery rate of 0.95 were implemented in the analysis. We classified the phenotypic trait into lactose intolerant (glucose level < 1.1 mmol/l), intermediate tolerance (1.1 mmol/l < glucose level < 1.7 mmol/l) and lactose tolerant (glucose level > 1.7 mmol/l), as implemented in previous studies (Arola 1994; Tishkoff et al. 2007;

Ranciaro et al. 2014). We applied the qtscore function in GenABEL and controlled for the population group the samples belonged to. Two additional GWAS runs were performed where we added the top SNPs (–13910 or rs6563275) as co-variates. Finally, to calculate how much a certain SNP explained the phenotype, a linear model estimation were used. We calculated the effect size of –13910 and rs6563275 in relation to the phenotypic residual variance and we used the adjusted R2 to infer the percentage of the phenotype explained by each SNP and both SNPs together.

### Selection Scans

Scans for signals of selection in the Fulani genomes were done using the integrated haplotype score (iHS) and Cross-population Extended Haplotype Homozygosity (XP-EHH) analyses within the R package REHH (Gautier et al. 2016). In both analyses the length of haplotype homozygosity was calculated with a maximum gap between two SNPs of 200,000 bp. We used the chimpanzee, bonobo and gorilla genomes to identify the ancestral allele and performed the selection analyses on 1,531,283 SNPs. Peaks were identified by averaging the −log10(p-value) every 10 SNPs and top 5 regions were inspected on Genome Browser to identify possible genes in the target region. Selection coefficient estimates were calculate using formula by Ohta and Kimura (Ohta and Kimura 1975).

## Results

### Fulani ancestry and admixture

We started by investigating the genetic affinities of the Fulani from Ziniaré in Burkina Faso using a set of comparative populations from Africa, Europe and Near East (Figure 1A, Table S1). The principal component analysis, PCA, (Figure 1B, S1) clusters the Fulani groups with other West Africans while displaying some genetic affinity to Eurasians. This prevalent West African component was also visible in population structure analysis (Figure 1C, S2), where the Fulani from Ziniaré in Burkina Faso have ancestry fractions of 74.5% West African, 21.4% European and 4.1% East African origin at K=3. We observe a similar genetic structure among all other Fulani groups in our dataset, except for the Fulani from Gambia. We notice that some individuals in this group display a higher European ancestry component than others, suggesting some degree of sub-structure in this population (Figure S2). This result might suggest recent additional admixture between certain Fulani groups from Gambia and West African neighbouring groups or alternatively, a shift in ethnic identity.

**Figure 1.**
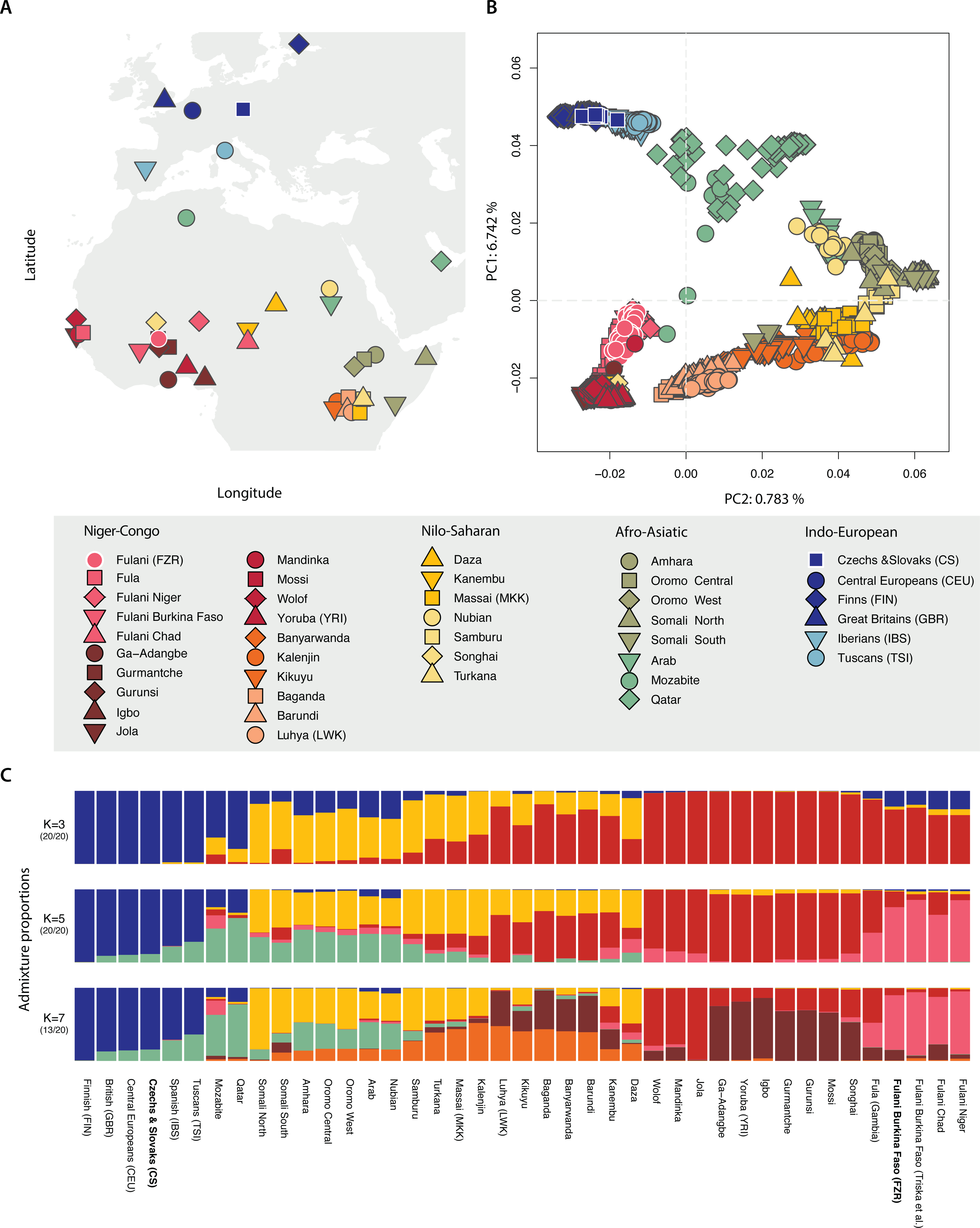
(A) Geographical location of the samples used in this study. (B) Principal component analysis and (C) Population averaged cluster analysis for K=3, 5 and 7 of the merged dataset of 1,355 individuals and 297,954 autosomal SNPs.

We inferred the time of admixture in Fulani genomes based on patterns of linkage disequilibrium decay (Patterson et al. 2012) and with a generation time of 29 years (Fenner 2005; Matsumura and Forster 2008), and found evidence for two admixture events between groups with West African and European ancestries (Table S2). The first admixture event is dated to 1828 years ago (95% CI: 1517-2138) between a parental population/s related to the West African ancestry groups in our dataset (Jola, Gurmantche, Gurunsi and Igbo) and a parental population carrying European ancestry (related to North-Western Europeans (CEU), Iberians (IBS), British (GBR), Tuscans (TSI), and Czech&Slovaks (CS) in our dataset). The second admixture event is dated to more recent times – 302 years ago (95% CI: 237–368) – and occurred between a West African group, with broadly similar ancestries compared to the first admixture event, and a European group. However, this European group is more related to present-day southwestern Europeans (Iberians (IBS) and Tuscans (TSI)).

In addition to the SNP typing we sequenced the LP region in intron 13 of the *MCMC* gene (upstream to *LCT* gene) in the Fulani, Czech and Slovak individuals, using Sanger sequencing. Of the known LP mutations in intron 13 of the *MCM6* gene, the Fulani from Ziniaré, Burkina Faso, only have the European LP T-13910 variant. We observed a T-13910 allele frequency of 48.0%, while the genome-wide European admixture fraction in the Fulani is 21.4% at K=3. The notable European admixture fraction in the Fulani coupled with the high frequencies of the LP T-13910 allele suggests the possibility of adaptive gene flow into the Fulani gene pool.

We reconstructed the local ancestry of the region surrounding the T-13910 allele and across the chromosome 2 for three Fulani samples (Fulani from Ziniaré, combined with West-Central African Fulani, and Fulani from Gambia), assuming either two or three ancestral sources: West African and European from the high density dataset A; and West African, European and North African from the lower density dataset B (Figures S3 and S4). The European genome proportions in the LP region were 0.519 and 0.491, for the two datasets respectively and in both cases all segments carrying the T-13910 allele were assigned to a European ancestry. The region extends for over 2Mb and contains 8 genes, including *LCT* and *MCM6* (Figure 2B) and haplotype lengths are similar in other Fulani groups in the dataset (Figure S3). For the dataset where North Africans were included as a parental population, the region near the LP variant departs 5.58 SD from the genome-wide average of European ancestry (μ=0.128, Figure S4). Looking in closer detail at the haplotype structure of this region, we observe that the haplotype carrying the mutation occurs at high frequency and show decreased diversity surrounding the T-13910 allele, compared to the alternative (ancestral) C-13910 allele (Figure S5, S6), indicating a strong selective sweep. Furthermore, in haplotype networks of the region, the haplotypes carrying the T-13910 allele in the Fulani cluster with European haplotypes (Figure S7). Our results therefore strongly support that the T-13910 LP allele occurs on a European haplotype background and was introduced into Fulani genomes by admixture rather that occurring as an independent convergent adaptation event.

**Figure 2.**
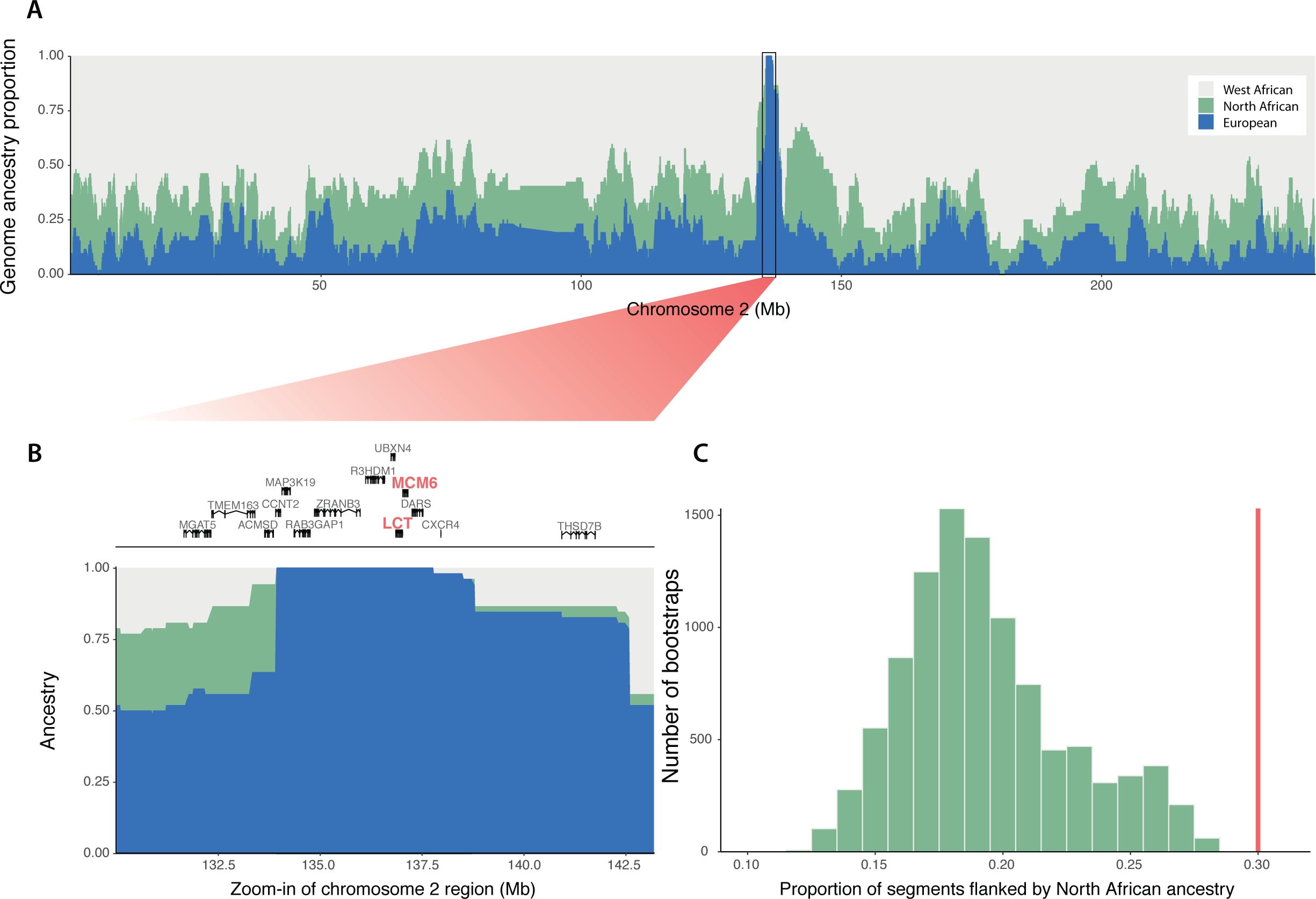
(A) Ancestry specific inference of chromosome 2 of haplotypes carrying allele T–13910 and (B) regional zoom-in. (C) Genome-wide distribution of randomly selected fragments being flanked by North-African-like segments over 10,000 bootstrap tests. The line in red represents the observed average proportion of European-like segments flanked by North-African-like segments in the Fulani from Burkina Faso.

To examine which particular source population was a likely candidate for this postulated European contact, we extracted all European-like segments across the Fulani genomes. We performed f3 outgroup analyses on the regions showing a European background (on the dataset with a separate North African component in the Fulani genomes – Extended Dataset B, Figure S8). The European-like segments showed the highest shared drift with Sardinians and French Basque populations, although based on the confidence intervals we could not specifically pinpoint any of the European groups included in the test. A previous study has reported a Mozabite-like (i.e. Berber-like) component in the Fulani from Burkina Faso and Niger (Triska et al. 2015), raising the possibility that the source population for the European admixture fraction (and LP mutation) could be of North African origin. This is difficult to observe in our clustering results since the Fulani form their own cluster (at K=4) before a North African component becomes visible (Figure 1C, S2). We therefore re-ran the clustering analysis with a supervised approach (Figure S9) and observed that the ancestry components of the Mozabite group could explain the non-West African genetic variation in the Fulani.

To further investigate the origin of the European ancestry segments in the Fulani, we analysed the flanking regions of European segments in their genomes. We observed a significant enrichment of North African ancestry in regions flanking European fragments. On average, European fragments in Fulani genomes are flanked by North African segments with a frequency of 0.302. To test for enrichment, we performed a bootstrapping test by randomly drawing fragments in the genome and recording their flanking regions (Figure 2C and Method section) and observed a highly significant association between European and North African segments in the Fulani genomes (p-value <1×10^−4^). These results suggest that it is unlikely that both ancestries would have been introduced by separate gene-flow events. To further test this, we simulated admixture scenarios (using genome-wide ancestry proportions of North Africa, Europe and West Africa in the Fulani genomes) and inferred the expected proportion of European haplotypes surrounded by North African ancestry in case of independent admixture events. If the European and North African segments were introduced by independent contact with a European and North African groups, respectively, we would expect on average that admixed segments would follow a random distribution across the genomes. In the 100 simulated populations we did not observe similar frequencies of European segments being surrounded by North African segments at the frequency we observe in the Fulani from Ziniaré, Burkina Faso (Figure S10, p-value <0.01), indicating that the two ancestries, at least in this Fulani population from Ziniaré, were not introduced by two separate events.

This scenario was further confirmed by testing specific demographic models using admixture graphs (Patterson et al. 2012). A model describing the Fulani as an admixed group between Mozabite and a West African group has a slightly lower Z-score (0.066) compared to a model where the Fulani result from admixture between a West African group and a western European group (CEU, 0.091) (Figure S11 A and B). However, when both Europeans and North Africans are included in the admixture graph models, a model that assumes that European ancestry is first admixed into North African ancestry and then introduced into the Fulani (Figure S11 C) is significant (Z-score = 0.926), whereas the model where Europeans directly mixed with West Africans to produce the Fulani is not significant (Figure S11 D).

### Lactase persistence in the Fulani

We established here that Fulani genomes acquired European admixture and the lactase persistence T-13910 allele by admixing with a North African population. Results from a Lactose Tolerence Test and Sanger sequeuncing on a larger group of Fulani, Czechs & Slovak individuals (see Method section) showed that carriers of the 13910*T allele (both TT–13910 and CT–13910 genotypes) have significantly higher glycemic levels than individuals homozygous for the –13910*C allele (Figure S12, S13, Table S3, S4). These results clearly associate the 13910*T allele with the LP phenotype and point to a dominant effect of the –13910*T allele in both Fulani and Czech & Slovak populations. Attempts to identify other regions in the genome associated with the ability to digest milk in adult life in a genome-wide setup have never been performed before, neither in the Fulani nor in any other group.

To investigate if other parts of Fulani genomes are involved in the ability to digest lactose we performed a Genome-Wide Association Scan (GWAS, Figure 3A, S14) for the glycemic measurement phenotype. This GWAS led to the identification of two regions, on chromosome 2 and chromosome 13 respectively, that clearly stand out. Even though none of the peaks reached the overly conservative Bonferroni multiple test correction threshold (due to small sample sizes and a large number of markers), the two prominent peaks on chromosome 2 and 14 clearly indicates an association with glucose levels in the bloodstream after ingestion of lactose. (Figure 3A). As expected the chromosome 2 peak overlaps with the region that contains the T-13910 mutation near the *LCT* gene (p-value = 3.17×10^−6^, Figure 3B). To test to what extent the –13910 SNP explain the phenotype, we calculated the effect size of the –13910 SNP based on a linear model. We observed that 35.1% of the residual variance can be explained by T-13910 allele (p-value = 3.709×10^−7^). Surprisingly, however, the region on chromosome 13 showed a slightly higher association with the phenotype in our GWAS analysis, with the highest association for the *rs6563275* SNP (p-value = 1.03×10^−6^, Figure 3C). This region does not contain any gene but it is located ~2.7 Mb upstream of the *SPRY2* gene (the nearest gene). The *rs6563275* SNP had an effect size of 38.7% (p-value = 6.62×10^−8^). For the rs6563275 and –13910 SNPs together, a combined effect size of 59.2% (p-value = 3.01×10^−12^) was estimated. The regions seams to act independent of each other and controlling for one SNP in the GWAS did not affect the other peak (Figure S15, S16). Also controlling for the top SNP in the two different regions seem to completely remove the association in the particular region, indicating that one SNP/haplotype per region is responsible for the associations (Figure S15B, S16B).

**Figure 3.**
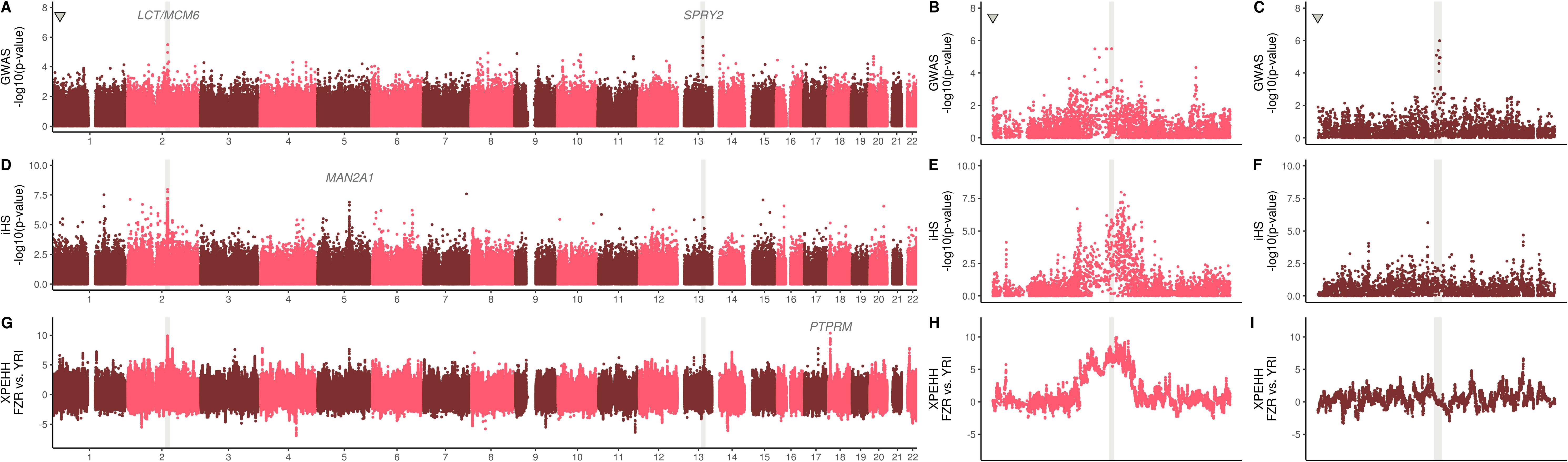
(A) P-values of the genome-wide association with the glycemic differentiation test after lactose ingestion. Triangular-shaped dot represents the Bonferroni p-value with alpha=0.05. (B, C) Zoom-ins of the chromosome 2 and 13 regions, respectively. (D) P-values of integrated haplotype scores (iHS) across the genome and (E, F) chromosome 2 and 13 regional zoom-ins. (G) FZR (Fulani, Burkina Faso) and YRI (Yoruba, Nigeria) cross-population extended heterozygosity haplotype (XP-EHH) across the genome and (H, I) chromosome 2 and 13 regional zoom-ins.

To test the impact of selection in Fulani genomes over the *LCT, SPRY2* and other regions across the whole genome, we calculated integrated haplotype scores (iHS) (Voight et al. 2006) and cross-population extended haplotype homozygosity (XP-EHH) (Sabeti et al. 2007) with the Yoruba as a comparative group (Figure 3D-I). Both tests showed clear signals of positive selection at the –13910 LP region on chromosome 2 in the Fulani. The LP region contained the highest peak for both scans (with 18.9 and 10.0 SD from genome-wide average, respectively). The XP-EHH results clearly showed the T-13910 allele as being selected in the Fulani compared to the Yoruba population (who does not carry any known LP variant) (Figure 3H). The region surrounding the *rs6563275* SNP on chromosome 13, however, did not display any signal of recent selection in our scans (Figure 3F, I). We calculated a selection coefficient for the –13910 LP region on chromosome 2 in the Fulani using Mozabite and CEU as parental populations, respectively (Figure S17). We found that a selection coefficient between 0.036 and 0.034 is necessary to explain the T-13910 allele frequency in the Fulani population, with the assumption of a constant allele frequency over time in the parental populations.

A number of other potential selection signals were observed across Fulani genomes (Table S5). A particular strong selection signal was observed on chromosome 18, where the XP-EHH test showed the second highest genome-wide region value (9.2 SD), comparable to that of the *MCM6/LCT* region. This signal seems to correspond to the *PTPRM* gene that encodes a tyrosine phosphatase enzyme highly expressed in adipose tissues and associated with HDL cholesterol levels, body weight and type 2 diabetes (Fox et al. 2007; Kathiresan et al. 2007; Murea et al. 2011). Furthermore, the iHS selection scan identified the region around the *MAN2A1* gene to be under selective pressure (p-value departing 17.0 SD from average). This gene encodes a glycosyl hydrolase found in the gut that functions in liberating *α*-glucose and *β*-glucose. Both these selection signals could represent additional indicators of dietary adaptation in the Fulani population.

## Discussion

The Fulani people are one of the most wide-spread pastoralist groups in the Sahel/Savannah belt, living (today) in a very large area that extends from the Fouta Djallon in Guinea to the Blue Nile in Ethiopia and Sudan. Even though an origin in the central Sahara has been suggested on archaeological grounds (Dupuy 1999), we found that the contemporary Fulani have a predominant West African genetic background combined with North African and European ancestry fractions (Figure 1B, S4, S9). These estimated genomic ancestry components, based on an in-depth genome analysis of a Fulani sample from Ziniaré, Burkina Faso, are comparable to those inferred in previously studied Fulani groups from other regions of Africa (Henn et al. 2012; Gurdasani et al. 2015; Triska et al. 2015). The sub-Saharan ancestry in Fulani clusters close to West African Niger-Congo speakers represented in our dataset by e.g. Wolof, Jola, Gurmanche, and Igbo (Figure 1B, S1, Table S2). The identification of the specific ancestry fragments flanking European-like segments, supervised admixture and demographic model predictions support the view that the European ancestry in Fulani genomes is coupled to their North African component (Figure 2C, S9-S11). These two genetic ancestries have been intertwined in the northwestern part of the African continent for the last 3,000 years (Fregel et al. 2017). Fregel and colleagues (2018) linked the diffusion of people across Gibraltar to Neolithic migrations and to the neolithic development in North Africa (Fregel et al. 2017). This trans-Gibraltar mixed ancestry was previously oberved in the Fulani mithocondrial gene-pool that link the Fulani to south-western Europe based on mtDNA haplogroups H1cb1 and U5b1b1b (Kulichova et al. 2017).

We inferred that the non-West African proportion in the Fulani were introduced through two admixture events (Table S2), dated to 1828 years ago (95% CI: 1517-2138) and 302 years ago (95% CI: 237–368). The oldest date compare well with previous dating efforts of the admixture event in the Fulani from Gambia (~1,800 years ago) (Busby et al. 2016; Busby et al. 2017), indicating a similar genetic history between the Fulani groups of Gambia and Burkina Faso. We hypothesize that the postulated first admixture between West African ancestors of the Fulani with an ancestral North African group/s possibly favoured, or even catalysed changes in their lifeways and consequently led the Fulani expansion throughout the Sahel/Savannah belt. This view is consistent with traces of pastoralism in the West African Savannah (northern Burkina Faso, in particular), starting around 2,000 years ago according to archaezoological data (Linseele 2013). The second admixture event dates to more recent times from a Southwestern European source (Table S2). This event can possibly be explained by either subsequent gene-flow between the Fulani and North Africans (who carries a considerate admixture proportions from Europeans due to trans-Gibraltar gene-flow); or by the colonial expansion into the African continent.

In the demographic model predictions where only one non-West African parental population is included (Figure S11 A and B), both European and North Africa can potentially explain the admixed part of the Fulani genetic composition. However, if both ancestries are present in the demographic model (Figure S11 C and D), only a North African ancestry population (mixed with a European population) can be a potential ancestor to the Fulani from Burkina Faso, whereas the model where Europeans directly mixed with West Africans to produce the Fulani is not significant. These results stress the importance of demographic context when identifying potential sources of admixture, when the sources have a similar genetic background.

The ability to digest milk during adulthood is a well-known case of recent selection in genomes of pastoralist and farming groups across the globe. The five independent mutations in intron 13 of the *MCM6* gene have been widely investigated and the association with expression of the *LCT* gene after the weaning period has been well established (Tishkoff et al. 2007; Gerbault et al. 2011; Liebert et al. 2017). The LP trait is associated with one of the most well-known signals of genetic adaptation to food-producing Neolithic lifestyles. High frequencies of the European-specific LP variant T-13910 are observed in Fulani groups across the Sahel/Savannah belt (Table S6). It is thought that the sustained expression of the *LCT* gene into adulthood, adds a dietary advantage in human populations who practice pastoralism for animal milk purposes. In our study the LP trait selection coefficient (s) estimates in the Fulani (Figure S17), 0.034 – 0.036, are comparable to previously calculated selection coeficients for LP in African populations; i.e. within East African groups it ranges between 0.035 and 0.077 (under a dominant model, (Tishkoff et al. 2007)), and 0.04-0.05 in Nama pastoralists of Southern Africa (Breton et al. 2014).

To date no publication has used a genome-wide approach to investigate whether other genomic regions are associated with the LP phenotype (Figure 3A-C, S14-S16). Here we confirmed an association between the previously identified chromosome 2 LP region on a genome-wide level. Additionally, we identified another signal associated with the ability to digest lactose (and generate glucose in the blood), on chromosome 13. We report here a strong association between glycemic levels (after lactose ingestion) and a region 2.7 Mb upstream of the *SPRY2* gene on chromosome 13. Previous GWAS studies have associated the *SPRY2* gene with adiposity and metabolism impairment (Kilpelainen et al. 2011), and with diabetes type 2 in Asian cohorts (Shu et al. 2010; Imamura et al. 2011; Mahajan et al. 2018). The importance of the association is possibly highlighted by a study that found that mice displayed hyperglycemia when the *SPRY2* gene is knocked down (Pappalardo et al. 2017), indicating that it is possible that the rate/extent of glucose formation is influenced by the *SPRY* gene. This gene have not previously been linked to the ability to digest lactose and possibly this region could be linked to an additional genetic variant that confers increased ability to digest lactose as adults. However, a more likely scenario is that the association we observe might not be because of LP trait itself, but rather due to the involvement of the genomic region in the subsequent steps of glycemic production. The latter hypothesis is supported by the fact that this region did not seem to have undergone positive selection in the Fulani, similar to the LP region on chromosome 2.

Genome-wide selection scans showed the chromosome 2 T-13910 region to be under strong selection, confirming that the European haplotype carrying the T-13910 mutation experienced adaptive gene-flow into the Fulani gene pool. Additional strong selection signals in the Fulani were found for genomic regions carrying the *MAN2A1* gene that encodes a glycosyl hydrolase and the *PTPRM* gene that encodes a tyrosine phosphatase expressed in the adipose tissues. These genes might represent other selection events in Fulani genomes to adapt to diets related to pastoralist lifeways. Higher consumption of sugars and fat contained in milk from domesticated animals might have triggered selective pressures in variants located within various genes leading to several dietary adaptations in the Fulani.

The complete history of the Fulani pastoralists remains to be uncovered, but through the genetic analyses performed in this study (based on the Fulani population from Ziniaré, Burkina Faso) we show that present-day Fulani genomic diversity developed from admixture between a West African group and a group/s that carried European and North African ancestry. The European LP variant was likely introduced through this admixture event, and was strongly selected in successive generations, in a similar way as the *TAS2R* gene family (Triska et al. 2015). Our results further showed that the LP region was not the only region that were under strong recent selective pressure in the Fulani ancestors, and several other selection signals points to dietary adaptations. It may well have been these and other similar selective advantages in the Fulani that contributed to their population expansion and long range spread across the Sahel/Savannah belt of Africa.

## Supplemental Data

Supplemental data contain six supplementary tables (S1-S6) as an accompanying excel file and 17 supplementary figures (S1-S17) as an accompanying pdf file.

## Acknowledgements

We are grateful to all subjects who participated in this research. We also would like to thank Torsten Günther, Paul Verdu, Jessica De Loma Olson and Yanjun Zan for helpful comments. We also thank Cecile Jolly for assisting with the Swedish Ethics permit application and submission. The genotyping was performed by the SNP&SEQ Technology Platform in Uppsala (www.genotyping.se). The facility is part of the National Genomics Infrastructure supported by the Swedish Research Council for Infrastructures and Science for Life Laboratory, Sweden. The SNP&SEQ Technology Platform is also supported by the Knut and Alice Wallenberg Foundation. The computations were performed at the Swedish National Infrastructure for Computing (SNIC-UPPMAX). The research project was supported by grants from the Swedish Research Council (no. 621-2014-5211) and the European Research Council (ERC – no. 759933) to CS and from the Grant Agency of the Czech Republic (grant 19-09352S-P505) to VČ. ESP was supported by Swiss National Science Foundation (grant 320030_159669).

## References

Alexander DH, Novembre J, Lange K (2009) Fast model-based estimation of ancestry in unrelated individuals. Genome Res 19:1655–1664

Arola H (1994) Diagnosis of hypolactasia and lactose malabsorption. Scand J Gastroenterol Suppl 202:26–35

Aulchenko YS, Ripke S, Isaacs A, van Duijn CM (2007) GenABEL: an R library for genome-wide association analysis. Bioinformatics 23:1294–1296

Auton A, Brooks LD, Durbin RM, Garrison EP, Kang HM, Korbel JO, Marchini JL, McCarthy S, McVean GA, Abecasis GR (2015) A global reference for human genetic variation. Nature 526:68–74

Ba AH, Daget J (1962) L’Empire peul du Macina. Mouton et Cie.

Bandelt HJ, Forster P, Rohl A (1999) Median-joining networks for inferring intraspecific phylogenies. Mol Biol Evol 16:37–48

Behr AA, Liu KZ, Liu-Fang G, Nakka P, Ramachandran S (2016) pong: fast analysis and visualization of latent clusters in population genetic data. Bioinformatics 32:2817–2823

Benoit M (1982) Nature peul du Yatenga. Remarques sur le pastoralisme en pays mossi. Travaux et Documents de l’ORSTOM 143

Bersaglieri T, Sabeti PC, Patterson N, Vanderploeg T, Schaffner SF, Drake JA, Rhodes M, Reich DE, Hirschhorn JN (2004) Genetic signatures of strong recent positive selection at the lactase gene. Am J Hum Genet 74:1111–1120

Boutrais J (1988) Des Peul en savanes humides. Développment pastoral dans l’ouest africain. Editions de l’ORSTOM, Paris

Breton G, Schlebusch CM, Lombard M, Sjodin P, Soodyall H, Jakobsson M (2014) Lactase persistence alleles reveal partial East african ancestry of southern african Khoe pastoralists. Curr Biol 24:852–858

Buckova J, Cerny V, Novelletto A (2013) Multiple and differentiated contributions to the male gene pool of pastoral and farmer populations of the African Sahel. Am J Phys Anthropol 151:10–21

Busby G, Christ R, Band G, Leffler E, Le QS, Rockett K, Kwiatkowski D, Spencer C (2017) Inferring adaptive gene-flow in recent African history. bioRxiv Jan 1:205252

Busby GB, Band G, Si Le Q, Jallow M, Bougama E, Mangano VD, Amenga-Etego LN, Enimil A, Apinjoh T, Ndila CM, Manjurano A, Nyirongo V, Doumba O, Rockett KA, Kwiatkowski DP, Spencer CC (2016) Admixture into and within sub-Saharan Africa. Elife 5

Cerezo M, Cerny V, Carracedo A, Salas A (2011) New insights into the Lake Chad Basin population structure revealed by high-throughput genotyping of mitochondrial DNA coding SNPs. PLoS One 6:e18682

Cerny V, Hajek M, Bromova M, Cmejla R, Diallo I, Brdicka R (2006) MtDNA of Fulani nomads and their genetic relationships to neighboring sedentary populations. Hum Biol 78:9–27

Cerny V, Kulichova I, Poloni ES, Nunes JM, Pereira L, Mayor A, Sanchez-Mazas A (2018) Genetic history of the African Sahelian populations. Hla 91:153–166

Chang CC, Chow CC, Tellier LC, Vattikuti S, Purcell SM, Lee JJ (2015) Second-generation PLINK: rising to the challenge of larger and richer datasets. Gigascience 4:7

Cizkova M, Hofmanova Z, Mokhtar MG, Janousek V, Diallo I, Munclinger P, Cerny V (2017) Alu insertion polymorphisms in the African Sahel and the origin of Fulani pastoralists. Ann Hum Biol 44:537–545

Coelho M, Sequeira F, Luiselli D, Beleza S, Rocha J (2009) On the edge of Bantu expansions: mtDNA, Y chromosome and lactase persistence genetic variation in southwestern Angola. BMC Evol Biol 9:80

de Bruijn M, van Dijk H (1995) Arid ways. Cultural understanding of insecurity in Fulbe society, central Mali. Thela publishers, Amsterdam

Delmet C (2000) Les Peuls nomades au Soudan. In: Diallo Y, Schlee G (eds) L’ethnicité peule dans des contextes nouveaux. Karthala, Paris, France, pp 191–206

Dupire M (1962) Peuls nomades, Étude descriptive des Wodaabé du Sahel nigérien. Karthala, Paris

Dupuy C (1999) Les apports de l’achéologie et de l’ethnologie a la connaisance de l’histoire ancienne des Peuls. In: Botte R, Boutrais J, Schmitz J (eds) Figures Peules. Karthala, Paris, pp 53–72

Durham W (1992) The evolution of adults lactose absortion. Coevolution Genes, Culture and Human Diversity. Standford University Press, Standford

Enattah NS, Jensen TG, Nielsen M, Lewinski R, Kuokkanen M, Rasinpera H, El-Shanti H, Seo JK, Alifrangis M, Khalil IF, Natah A, Ali A, Natah S, Comas D, Mehdi SQ, Groop L, Vestergaard EM, Imtiaz F, Rashed MS, Meyer B, Troelsen J, Peltonen L (2008) Independent introduction of two lactase-persistence alleles into human populations reflects different history of adaptation to milk culture. Am J Hum Genet 82:57–72

Fay C (1997) Les derniers seront les premiers: peuplement et pouvoirs mandingues et peuls au Maasina (Mali). In: De Bruijn M, Van Dijk H (eds) Peuls et Mandingues: Dialectique des Constructions Identitaires. Kathala, Paris, pp 165–191

Fenner JN (2005) Cross-cultural estimation of the human generation interval for use in genetics-based population divergence studies. Am J Phys Anthropol 128:415–423

Fox CS, Heard-Costa N, Cupples LA, Dupuis J, Vasan RS, Atwood LD (2007) Genome-wide association to body mass index and waist circumference: the Framingham Heart Study 100K project. BMC Med Genet 8 Suppl 1:S18

Fregel R, Mendez FL, Bokbot Y, Martin-Socas D, Camalich-Massieu MD, Avila-Arcos MC, Underhill PA, Shapiro B, Wojcik GL, Rasmussen M, Soares AER, Kapp J, Sockell A, Rodriguez-Santos FJ, Mikdad A, Santana J, Trujillo-Mederos A, Bustamante CD (2017) Neolithization of North Africa involved the migration of people from both the Levant and Europe. bioRxiv 191569

Gautier M, Klassmann A, Vitalis R (2016) rehh 2.0: a reimplementation of the R package rehh to detect positive selection from haplotype structure. Mol Ecol Resour 17:78–90

Gerbault P, Liebert A, Itan Y, Powell A, Currat M, Burger J, Swallow DM, Thomas MG (2011) Evolution of lactase persistence: an example of human niche construction. Philos Trans R Soc Lond B Biol Sci 366:863–877

Greenberg JH (1963) The languages of Africa. Indiana University Press, Bloomington, Indiana

Gurdasani D, Carstensen T, Tekola-Ayele F, Pagani L, Tachmazidou I, Hatzikotoulas K, Karthikeyan S, et al. (2015) The African Genome Variation Project shapes medical genetics in Africa. Nature 517:327–332

Haber M, Mezzavilla M, Bergstrom A, Prado-Martinez J, Hallast P, Saif-Ali R, Al-Habori M, Dedoussis G, Zeggini E, Blue-Smith J, Wells RS, Xue Y, Zalloua PA, Tyler-Smith C (2016) Chad Genetic Diversity Reveals an African History Marked by Multiple Holocene Eurasian Migrations. Am J Hum Genet

Henn BM, Botigue LR, Gravel S, Wang W, Brisbin A, Byrnes JK, Fadhlaoui-Zid K, Zalloua PA, Moreno-Estrada A, Bertranpetit J, Bustamante CD, Comas D (2012) Genomic ancestry of North Africans supports back-to-Africa migrations. PLoS Genet 8:e1002397

Hollox EJ, Poulter M, Zvarik M, Ferak V, Krause A, Jenkins T, Saha N, Kozlov AI, Swallow DM (2001) Lactase haplotype diversity in the Old World. Am J Hum Genet 68:160–172

Imamura M, Iwata M, Maegawa H, Watada H, Hirose H, Tanaka Y, Tobe K, Kaku K, Kashiwagi A, Kawamori R, Nakamura Y, Maeda S (2011) Genetic variants at CDC123/CAMK1D and SPRY2 are associated with susceptibility to type 2 diabetes in the Japanese population. Diabetologia 54:3071–3077

Itan Y, Jones BL, Ingram CJ, Swallow DM, Thomas MG (2010) A worldwide correlation of lactase persistence phenotype and genotypes. BMC Evol Biol 10:36

Johnston HAS (1967) The Fulani Empire of Sokoto. Oxford University Press, Oxford

Kathiresan S, Manning AK, Demissie S, D’Agostino RB, Surti A, Guiducci C, Gianniny L, Burtt NP, Melander O, Orho-Melander M, Arnett DK, Peloso GM, Ordovas JM, Cupples LA (2007) A genome-wide association study for blood lipid phenotypes in the Framingham Heart Study. BMC Med Genet 8 Suppl 1:S17

Kilpelainen TO, Zillikens MC, Stancakova A, Finucane FM, Ried JS, Langenberg C, Zhang W, et al. (2011) Genetic variation near IRS1 associates with reduced adiposity and an impaired metabolic profile. Nat Genet 43:753–760

Kulichova I, Fernandes V, Deme A, Novackova J, Stenzl V, Novelletto A, Pereira L, Cerny V (2017) Internal diversification of non-Sub-Saharan haplogroups in Sahelian populations and the spread of pastoralism beyond the Sahara. Am J Phys Anthropol 164:424–434

Lam AM (2001) De l’origine égyptienne des Peuls. Editions Présence Africaine

Liebert A, Lopez S, Jones BL, Montalva N, Gerbault P, Lau W, Thomas MG, Bradman N, Maniatis N, Swallow DM (2017) World-wide distributions of lactase persistence alleles and the complex effects of recombination and selection. Hum Genet 136:1445–1453

Linseele V (2013) From the first stock keepers to specialised pastoralists in the West African Savannah. In: Bollig M, Schnegg M, Wotzka H-P (eds) Pastoralism in Africa: Past, Present and Future. Berghan Books, New York

Lokki AI, Jarvela I, Israelsson E, Maiga B, Troye-Blomberg M, Dolo A, Doumbo OK, Meri S, Holmberg V (2011) Lactase persistence genotypes and malaria susceptibility in Fulani of Mali. Malar J 10:9

Macholdt E, Lede V, Barbieri C, Mpoloka SW, Chen H, Slatkin M, Pakendorf B, Stoneking M (2014) Tracing pastoralist migrations to southern Africa with lactase persistence alleles. Curr Biol 24:875–879

Mahajan A, Taliun D, Thurner M, Robertson NR, Torres JM, Rayner NW, Payne AJ, et al. (2018) Fine-mapping type 2 diabetes loci to single-variant resolution using high-density imputation and islet-specific epigenome maps. Nat Genet 50:1505–1513

Maples BK, Gravel S, Kenny EE, Bustamante CD (2013) RFMix: a discriminative modeling approach for rapid and robust local-ancestry inference. Am J Hum Genet 93:278–288

Matsumura S, Forster P (2008) Generation time and effective population size in Polar Eskimos. Proc Biol Sci 275:1501–1508

McIntosh R, McIntosh SK, Bocoum H (2016) The Search for Takrur: Archaeological Excavations and Reconnaissance along the Middle Senegal Valley. The Yale Peabody Museum

Mulcare CA, Weale ME, Jones AL, Connell B, Zeitlyn D, Tarekegn A, Swallow DM, Bradman N, Thomas MG (2004) The T allele of a single-nucleotide polymorphism 13.9 kb upstream of the lactase gene (LCT) (C-13.9kbT) does not predict or cause the lactase-persistence phenotype in Africans. Am J Hum Genet 74:1102–1110

Murea M, Lu L, Ma L, Hicks PJ, Divers J, McDonough CW, Langefeld CD, Bowden DW, Freedman BI (2011) Genome-wide association scan for survival on dialysis in African-Americans with type 2 diabetes. Am J Nephrol 33:502–509

Newman JL (1995) The peopling of Africa. Yale University Press, New Haven, CT

O’Connell J, Gurdasani D, Delaneau O, Pirastu N, Ulivi S, Cocca M, Traglia M, et al. (2014) A general approach for haplotype phasing across the full spectrum of relatedness. PLoS Genet 10:e1004234

Ohta T, Kimura M (1975) The effect of selected linked locus on heterozygosity of neutral alleles (the hitch-hiking effect). Genetical research 25:313–325

Pappalardo Z, Gambhir Chopra D, Hennings TG, Richards H, Choe J, Yang K, Baeyens L, Ang K, Chen S, Arkin M, German MS, McManus MT, Ku GM (2017) A Whole-Genome RNA Interference Screen Reveals a Role for Spry2 in Insulin Transcription and the Unfolded Protein Response. Diabetes 66:1703–1712

Patterson N, Moorjani P, Luo Y, Mallick S, Rohland N, Zhan Y, Genschoreck T, Webster T, Reich D (2012) Ancient admixture in human history. Genetics 192:1065–1093

Patterson N, Price AL, Reich D (2006) Population structure and eigenanalysis. PLoS Genet 2:e190

Podgorna E, Diallo I, Vangenot C, Sanchez-Mazas A, Sabbagh A, Cerny V, Poloni ES (2015) Variation in NAT2 acetylation phenotypes is associated with differences in food-producing subsistence modes and ecoregions in Africa. BMC Evol Biol 15:263

Polzin T, Daneschmand SV (2003) On Steiner trees and minimum spanning trees in hypergraphs. Operations Res Lett 31:12–20

Price AL, Patterson NJ, Plenge RM, Weinblatt ME, Shadick NA, Reich D (2006) Principal components analysis corrects for stratification in genome-wide association studies. Nat Genet 38:904–909

Priehodova E, Abdelsawy A, Heyer E, Cerny V (2014) Lactase persistence variants in Arabia and in the African Arabs. Hum Biol 86:7–18

Priehodova E, Austerlitz F, Cizkova M, Mokhtar MG, Poloni ES, Cerny V (2017) The historical spread of Arabian Pastoralists to the eastern African Sahel evidenced by the lactase persistence –13,915*G allele and mitochondrial DNA. Am J Hum Biol 29

Ranciaro A, Campbell MC, Hirbo JB, Ko WY, Froment A, Anagnostou P, Kotze MJ, Ibrahim M, Nyambo T, Omar SA, Tishkoff SA (2014) Genetic origins of lactase persistence and the spread of pastoralism in Africa. Am J Hum Genet 94:496–510

Riesman R (1974) Société et liberté chez les Peul Djelgôbé de Haute Volta. La Haye, Mouton, Paris

Sabeti PC, Varilly P, Fry B, Lohmueller J, Hostetter E, Cotsapas C, Xie X, et al. (2007) Genome-wide detection and characterization of positive selection in human populations. Nature 449:913–918

Schlebusch CM (2019) Population migration and adaptation during the African Holocene: A genetic perspective. In: Sahle Y, Reyes-Centeno H, Bentz C (eds) Modern Human Origins and Dispersal. Kerns Verlag, Tuebingen, Germany

Schlebusch CM, Jakobsson M (2018) Tales of Human Migration, Admixture, and Selection in Africa. Annu Rev Genomics Hum Genet 19:405–428

Schlebusch CM, Sjodin P, Skoglund P, Jakobsson M (2012a) Stronger signal of recent selection for lactase persistence in Maasai than in Europeans. Eur J Hum Genet

Schlebusch CM, Skoglund P, Sjodin P, Gattepaille LM, Hernandez D, Jay F, Li S, De Jongh M, Singleton A, Blum MG, Soodyall H, Jakobsson M (2012b) Genomic Variation in Seven Khoe-San Groups Reveals Adaptation and Complex African History. Science 338:374–379

Schultz EA (1984) From Pagan to Pullo: Ethnic Identity Change in Northern Cameroon. Africa 54:33–65

Shu XO, Long J, Cai Q, Qi L, Xiang YB, Cho YS, Tai ES, et al. (2010) Identification of new genetic risk variants for type 2 diabetes. PLoS Genet 6:e1001127

Stenning DJ (1957) Transhumance, Migratory Drift, Migration; Patterns of Pastoral Fulani Nomadism. The Journal of the Royal Anthropological Institute of Great Britain and Ireland 87:57–73

Swallow DM (2003) Genetics of lactase persistence and lactose intolerance. Annu Rev Genet 37:197–219

Tishkoff SA, Reed FA, Ranciaro A, Voight BF, Babbitt CC, Silverman JS, Powell K, Mortensen HM, Hirbo JB, Osman M, Ibrahim M, Omar SA, Lema G, Nyambo TB, Ghori J, Bumpstead S, Pritchard JK, Wray GA, Deloukas P (2007) Convergent adaptation of human lactase persistence in Africa and Europe. Nat Genet 39:31–40

Triska P, Soares P, Patin E, Fernandes V, Cerny V, Pereira L (2015) Extensive Admixture and Selective Pressure Across the Sahel Belt. Genome Biol Evol 7:3484–3495

Voight BF, Kudaravalli S, Wen X, Pritchard JK (2006) A map of recent positive selection in the human genome. PLoS Biol 4:e72

