## Supplementary Figures for "Population history and genetic adaptation of the Fulani nomads: Inferences from genome-wide data and the lactase persistence trait"

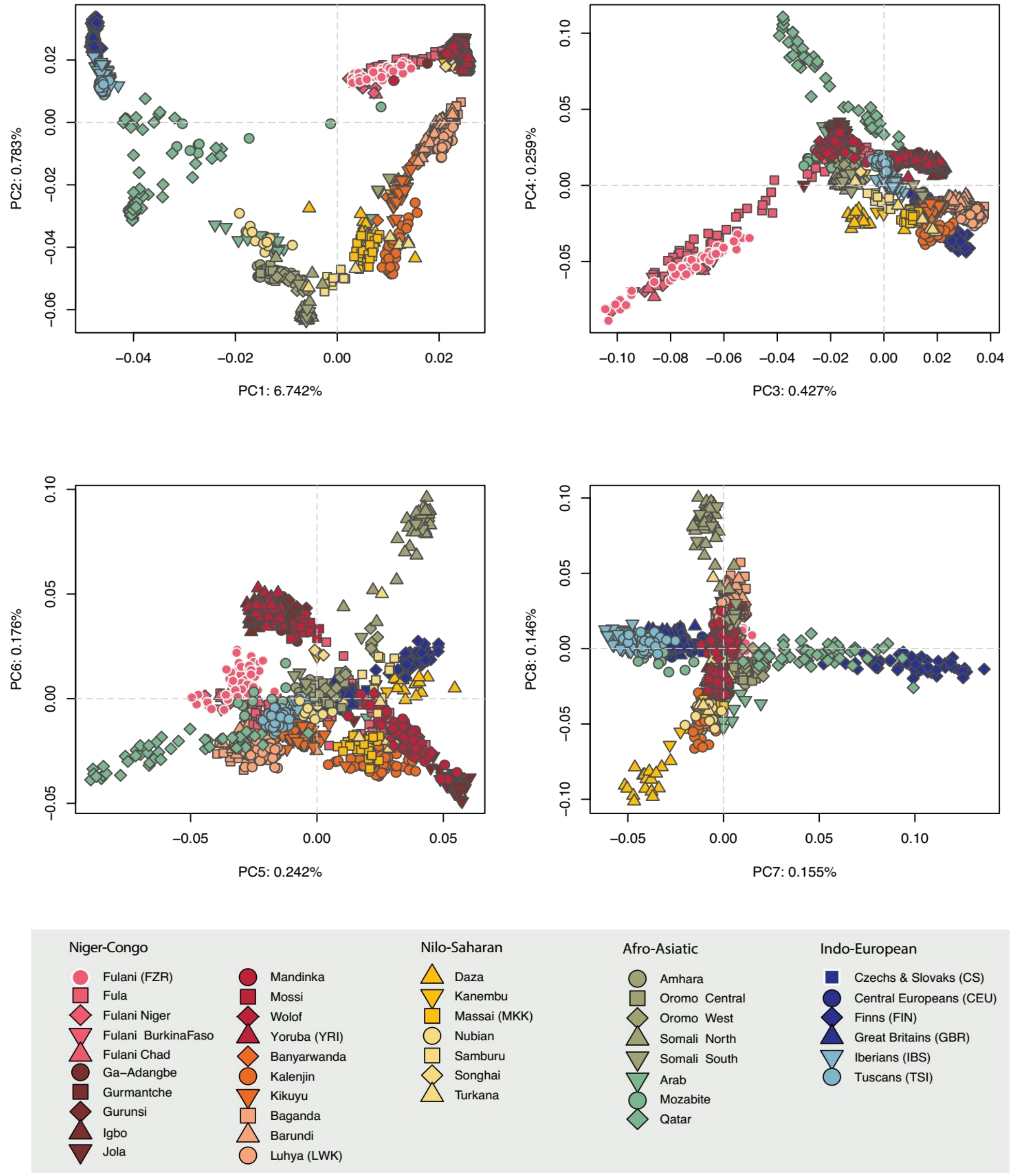


***Figure S1 - Principal Component Analysis (PCA) plots - PC 1 to 8 - based on 1,355 individuals with 297,954 autosomal SNPs (dataset B)****. The uppermost left plot is shown in Figure 1B.*

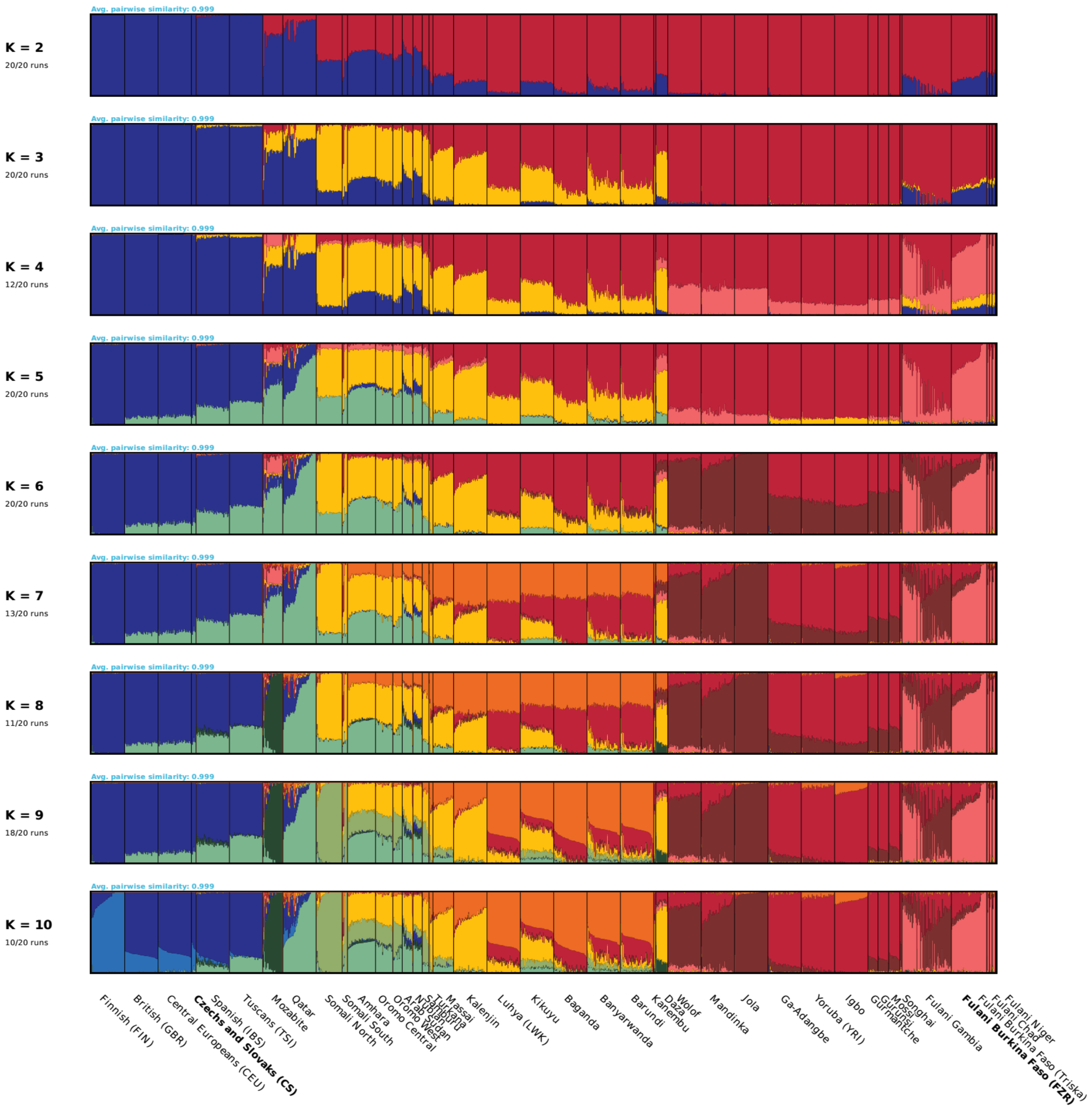


***Figure S2 - Admixture clustering analysis of dataset B.*** *The analyses are based on 1,355 individuals with 297,954 autosomal SNPs (dataset B).*

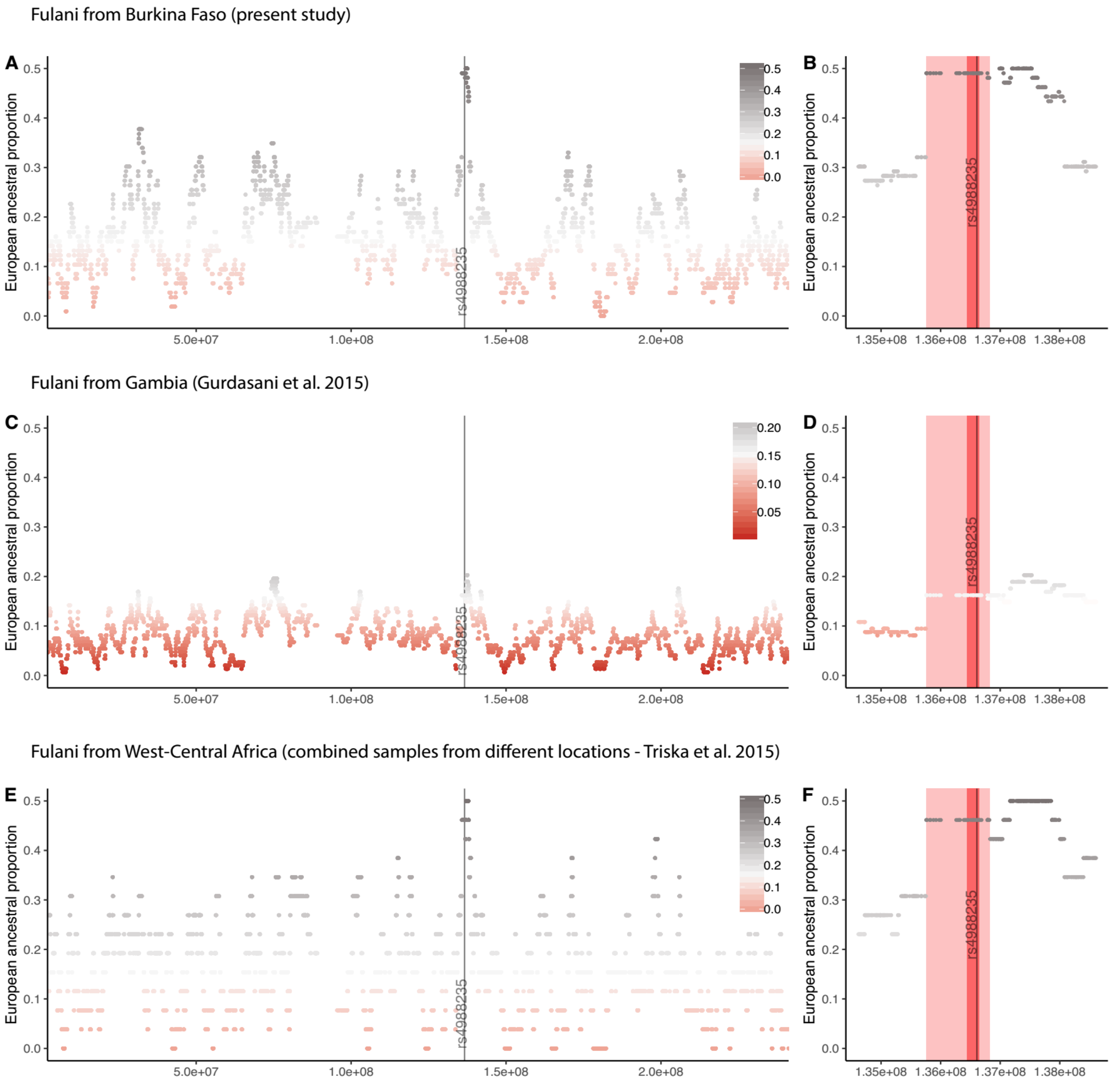


***Figure S3 - European ancestry proportion across chromosome 2 (A, C and E) and near the C/T–13910 (B, D and F) in three Fulani groups.*** *The analyses were done on the low SNP density dataset B for the Fulani from Burkina Faso (present study), Fulani from Gambia (Gurdasani et al. 2015), and combined Fulani from West-Central Africa (Triska et al. 2015).*

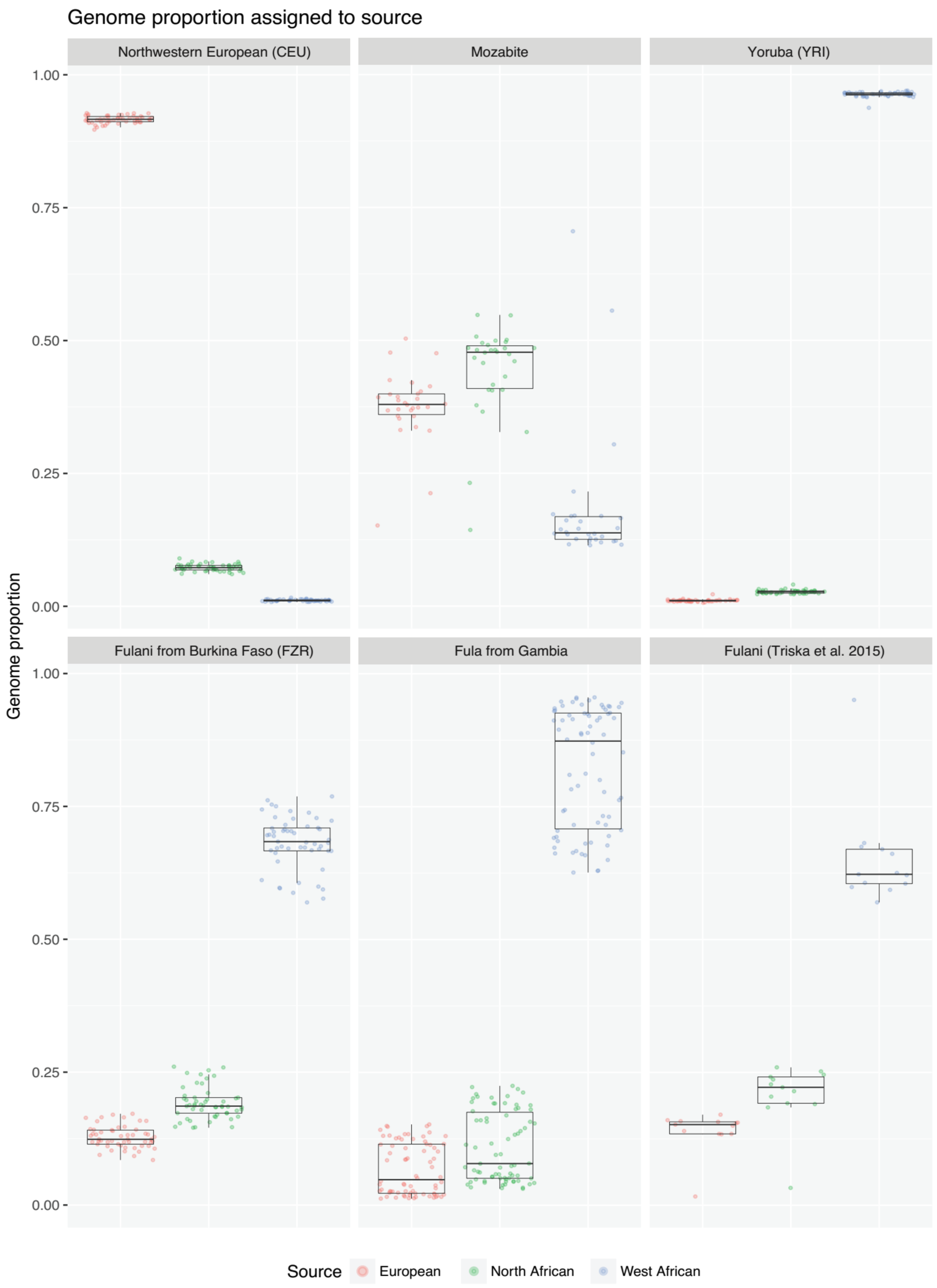


***Figure S4 - Genome ancestry proportion for the Fulani from Burkina Faso (FZR), Fulani from West-Central Africa, Fula from Gambia and the parental sources (CEU, Mozabite and Yoruba).*** *The results were obtained using in RFMix on 297,954 autosomal SNPs (dataset B).*

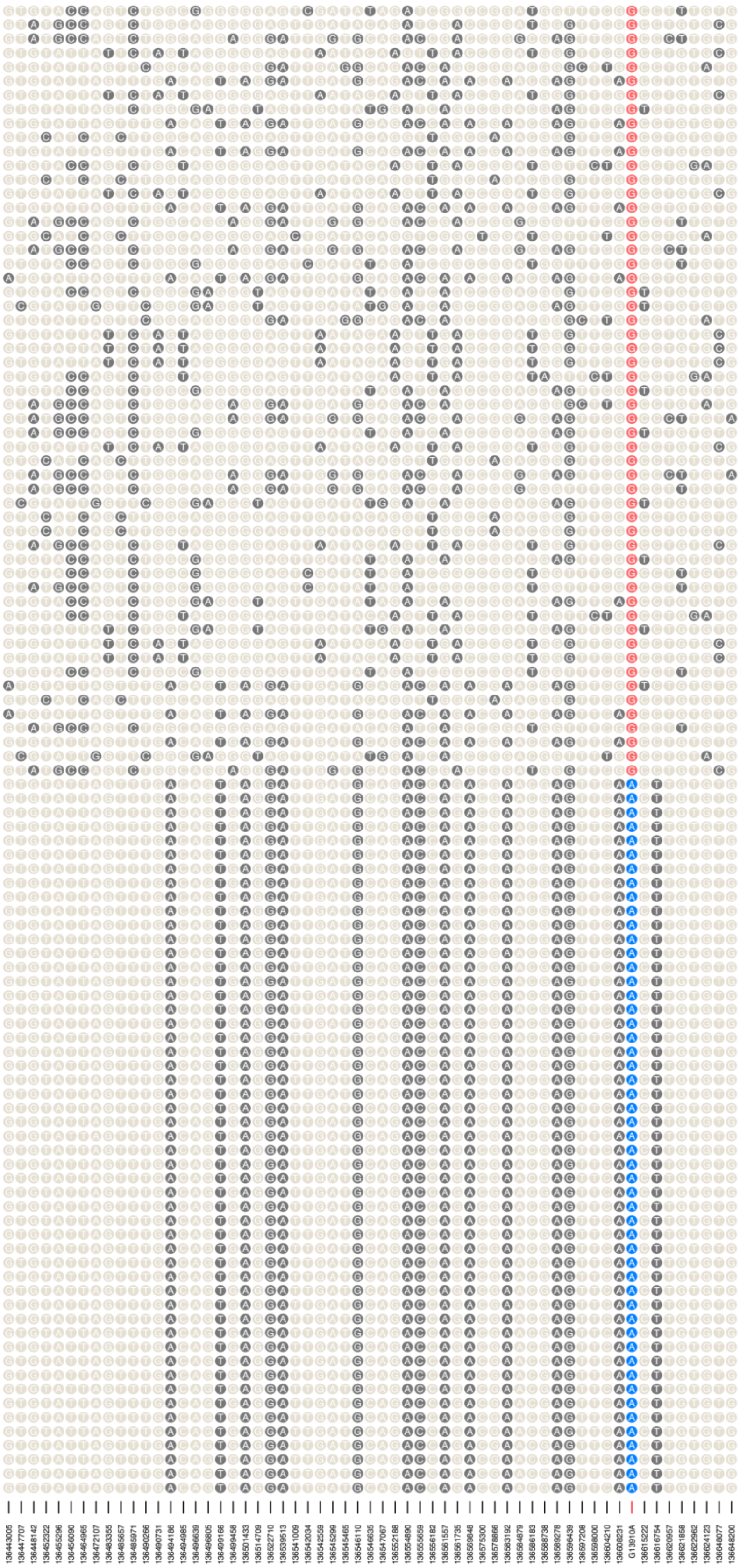


***Figure S5 - Haplotype background around the lactase persistence 13910 region.*** *The displayed region was based on RFMix assignment of the T-13910 window. The haplotypes are displayed based on the complementary strand.*

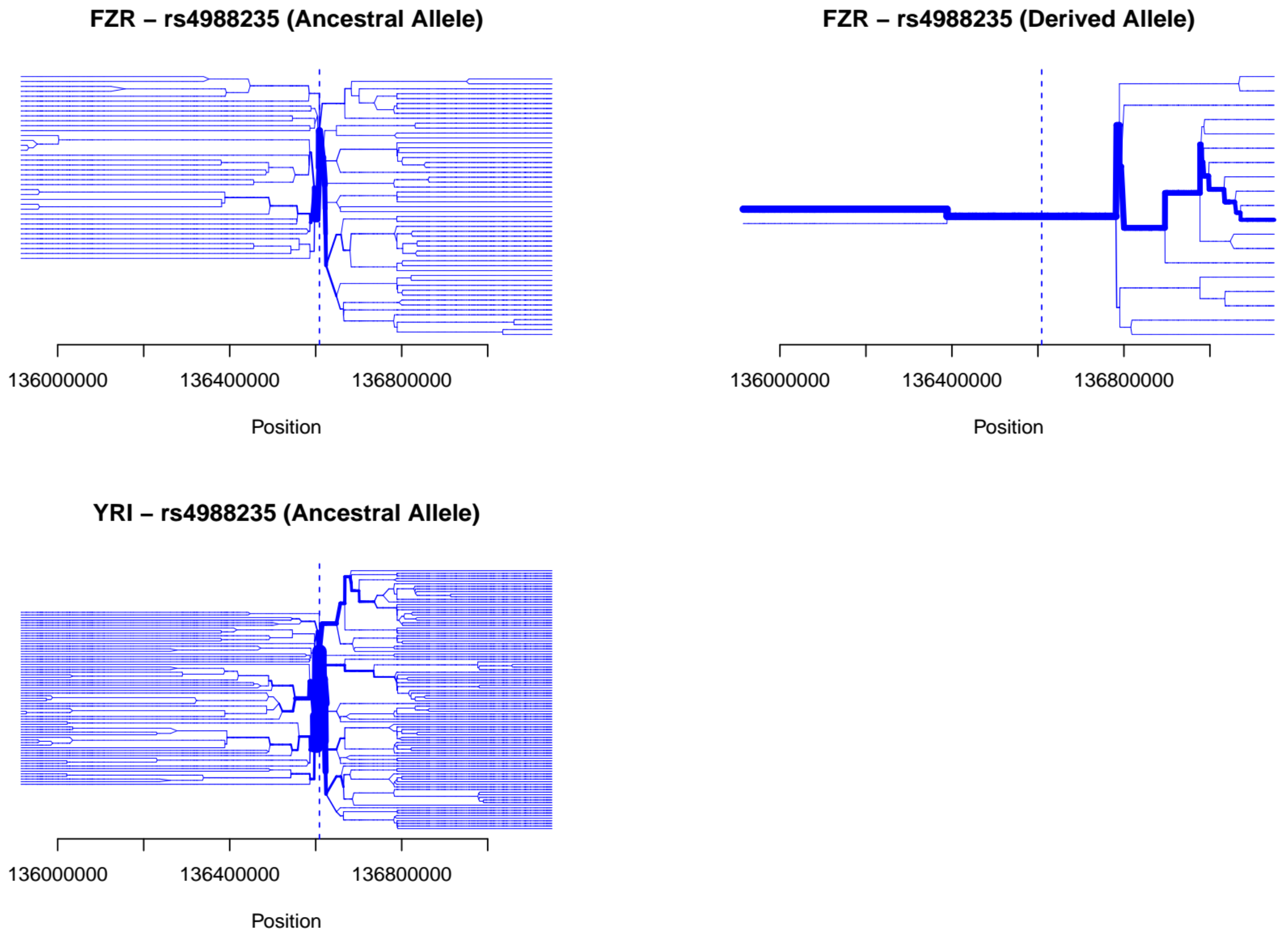


***Figure S6 - Bifurcation plots based on the haplotype structure around -13910 for the ancestral (C) and derived (T) allele in the Fulani from Burkina Faso and Yoruba (who only carry the ancestral allele).***

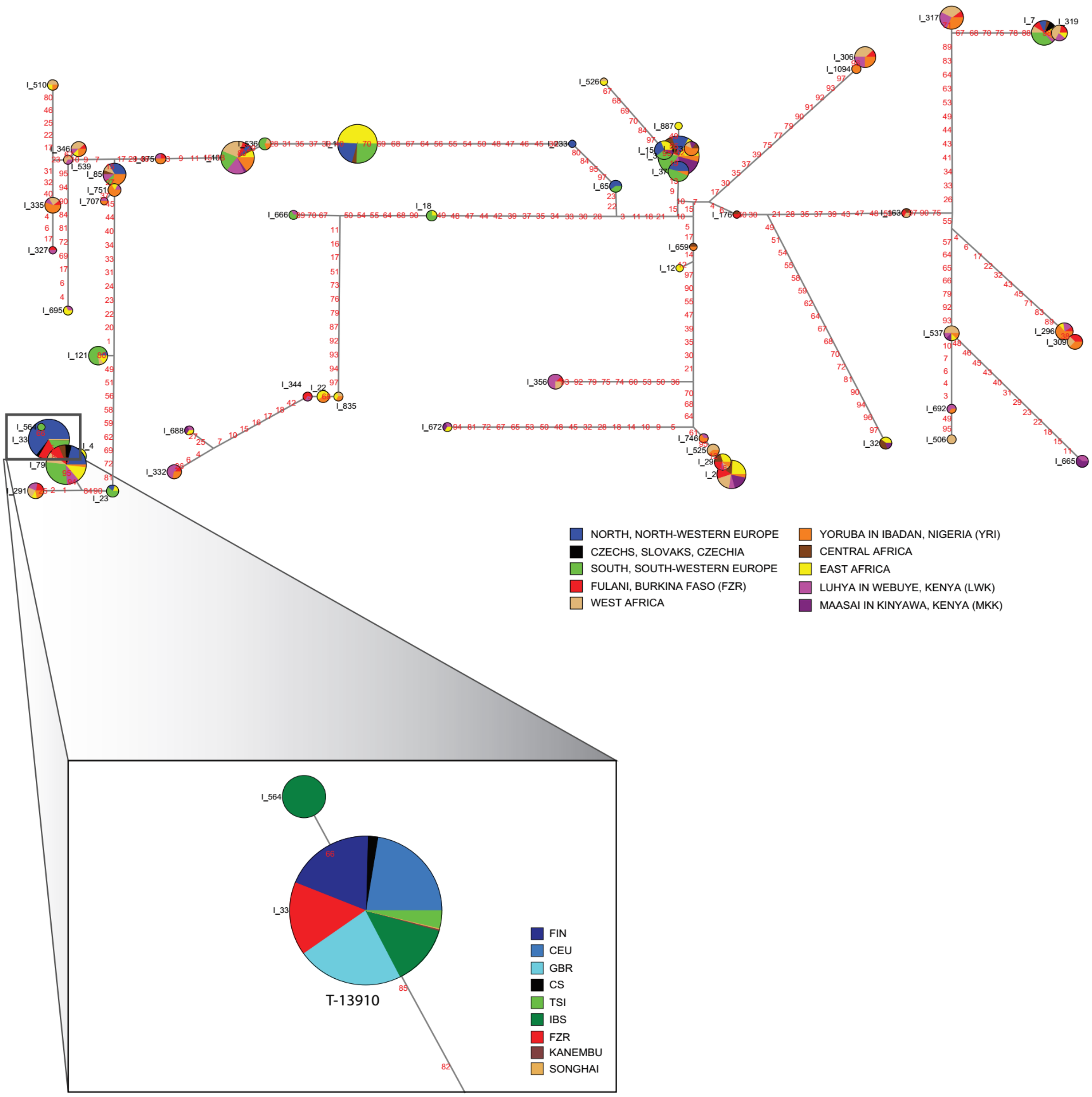


***Figure S7 - A: Network analysis based on 98 SNPs haplotype in a 0.2 Mb long fragment around –13910 for European and African population samples from dataset A.*** *For simplification, some population samples were merged: NORTH, NORTH-WESTERN EUROPE – CEU, FIN, GBR; SOUTH, SOUTH-WESTERN EUROPE – IBS, TSI; WEST AFRICA, CENTRAL AFRICA, EAST AFRICA. Zoom-in on the network branch is defined by –13910*T.*

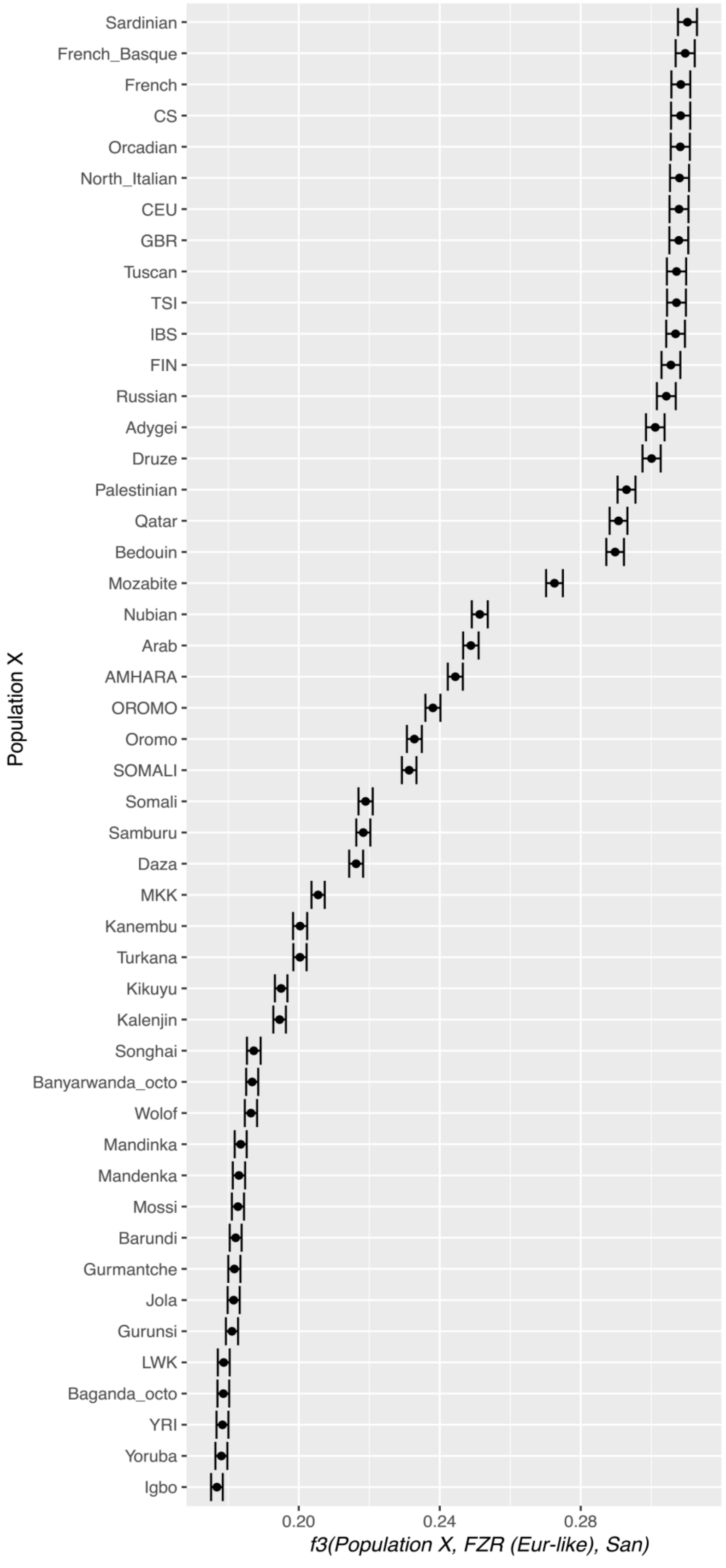


***Figure S8 - f3-outgroup(Population X, European-like FZR, San) results.*** *Only the European-like ancestry fragments of Fulani were analyzed.*

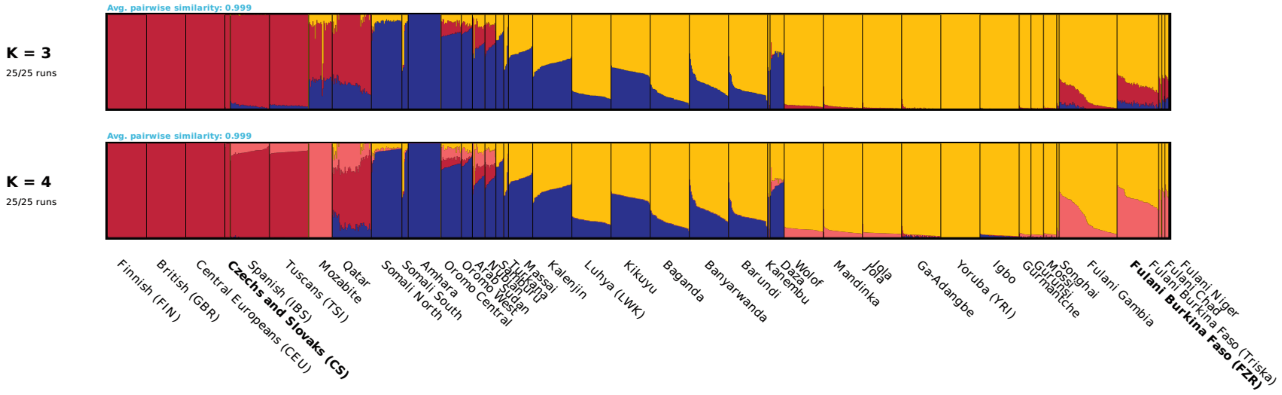


***Figure S9 - Supervised admixture clustering analysis of dataset B.*** *The analyses are based on 1,355 individuals with 297,954 autosomal SNPs (dataset B). For K=3 we have supervised with CEU, Amhara and Yoruba populations; at K=4 we added Mozabite to the reference groups of K=3.*

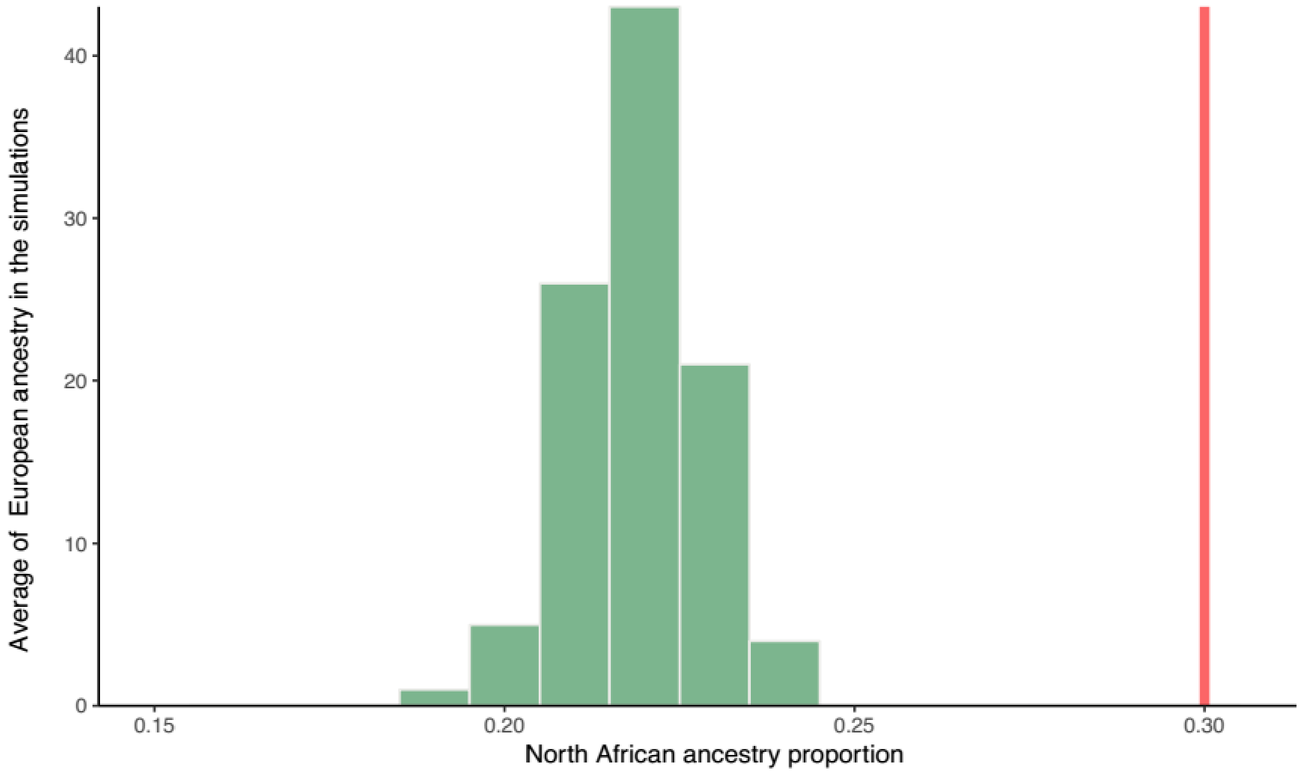


***Figure S10 - Average genome-wide proportion of European fragments being flanked by North African ancestry regions in 100 simulated populations.*** *The simulations assumed that the two different non-West Africa ancestries were introduced in the Fulani genomes by two distinct admixture events. The Fulani genome-wide proportions of European, North African and West African ancestries, were used as starting inputs in the simulated populations. The line in red represents the observed average proportion of European-like segments flanked by North-African-like segments in the Fulani from Burkina Faso.*

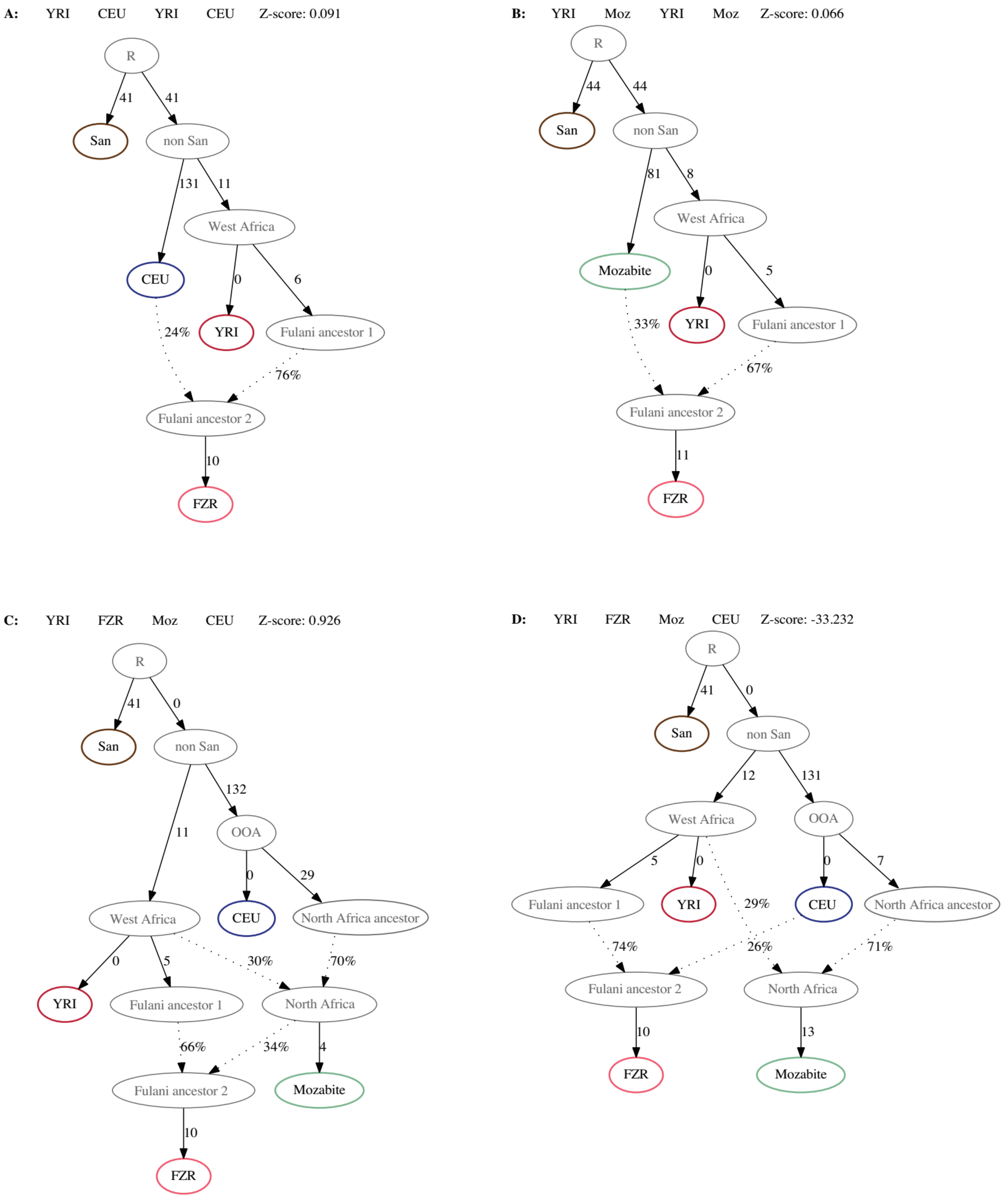


***Figure S11 - Demographic model testing using qpGraph.*** *Colored outlines indicate a sampled population and ancestral nodes are colored in gray. "R" stand for Root and "OOA" for the Out-of-Africa.* ***A****: Using San, CEU, Yoruba and Fulani as real populations.* ***B****: San, Mozabite, Yoruba (YRI) and Fulani.* ***C*** *and* ***D****: San, CEU, Mozabite, Yoruba and Fulani. When incorporating both CEU and Mozabite in the model the Fulani genetic composition can only be explained by admixture of a European group through contact with North African ancestry (model C). In model D, direct admixture from a European group does not lead to a significant explanation of Fulani ancestry.*

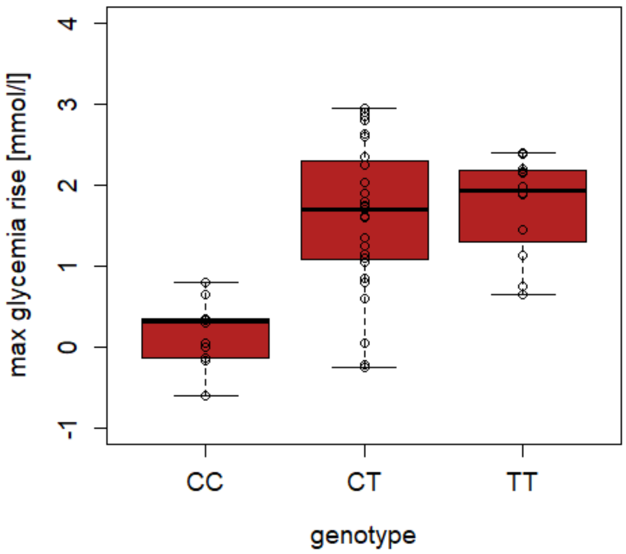


***Figure S12 - Maximal glycemia for non-LP genotype CC–13910 and LP-genotypes CT–13910 and TT–13910 in the 53 Fulani samples of Ziniaré (FZR), Burkina Faso.*** *Carriers of LP mutation –13910*T (both TT–13910 and CT–13910 genotypes) have significantly higher blood glucose levels after consummation of 50 g of lactose than those carrying only the –13910*C variant. These results are confirmed by Wilcoxon signed-rank test which shows that phenotype of bearers of LP mutation –13910*T differs from persons without the mutation – see Table S3.*

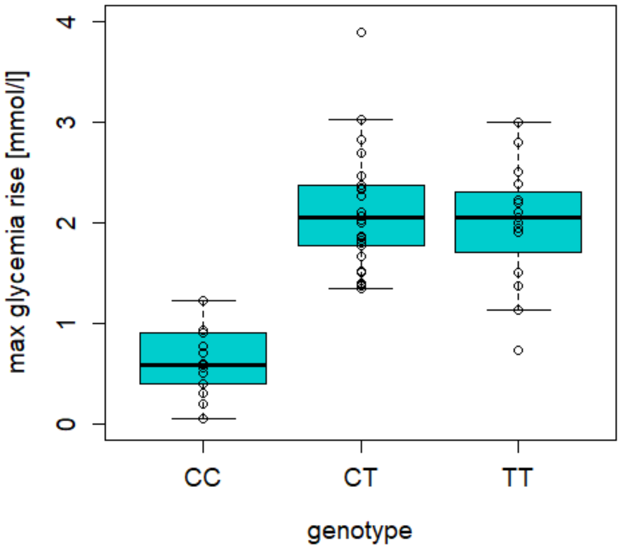


***Figure S13 - Maximal glycemia for non-LP genotype CC–13910 and LP-genotypes CT–13910 and TT–13910 in Czechs and Slovaks (n=63).*** *In a dataset of Czech/Slovak samples (n=63), the sequencing of 359 bp fragment intron 13 of the MCM6 gene confirms the presence of only one LP mutation –13910*T, whose frequency is 0.49. As in the phenotyped sample of Fulani from Burkina Faso, the Czech/Slovak sample with LP mutation –13910*T has higher blood glucose level after consummation of 50 g of lactose than persons with –13910*C. These results are confirmed by Wilcoxon signed-rank test which shows that phenotype of bearers of LP mutation –13910*T differs from persons without the mutation – see Table S4. (Due to DNA condition only a subset of 7 from these samples were used for SNP genotyping and further analyses).*

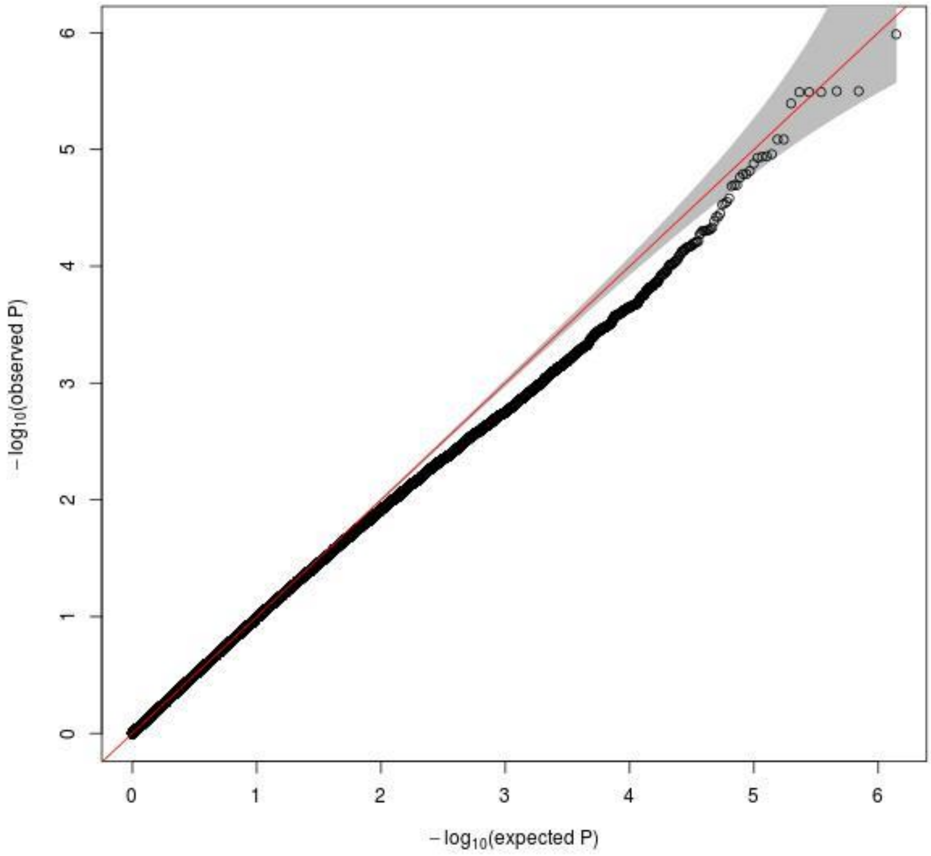


*Figure S14 - Quantile-quantile (Q-Q) plot of the Genome-Wide Association Study (GWAS).*

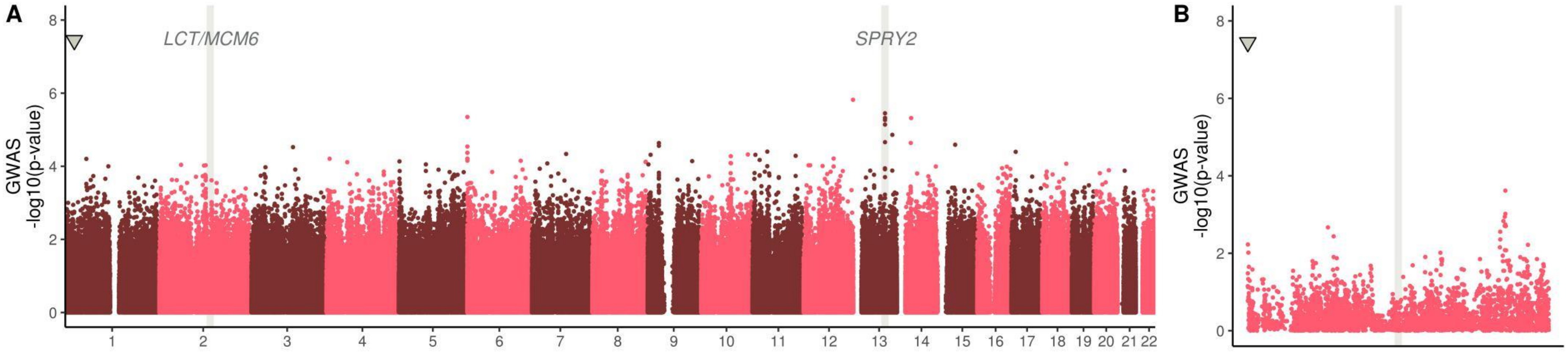


***Figure S15 -P-values of the genome-wide association with the glycemic differentiation test after lactose ingestion conditioned on group and -13910 SNP as co-variate. A:*** *Across the genome,* ***B:*** *LCT region zoom-in.*

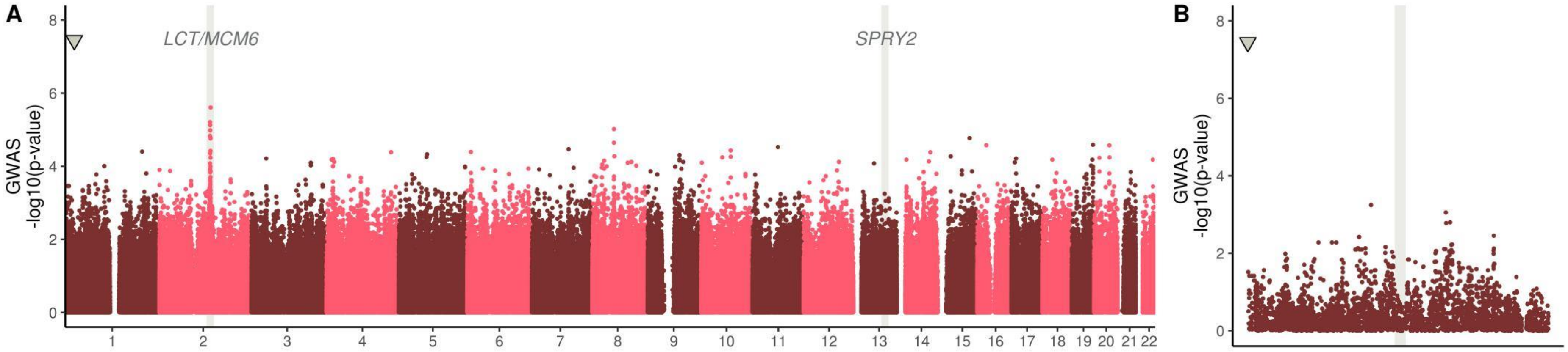


***Figure S16 - P-values of the genome-wide association with the glycemic differentiation test after lactose ingestion conditioned on group and rs6563275 SNP as co-variate. A:*** *Across the genome,* ***B:*** *SPRY2 region zoom-in.*

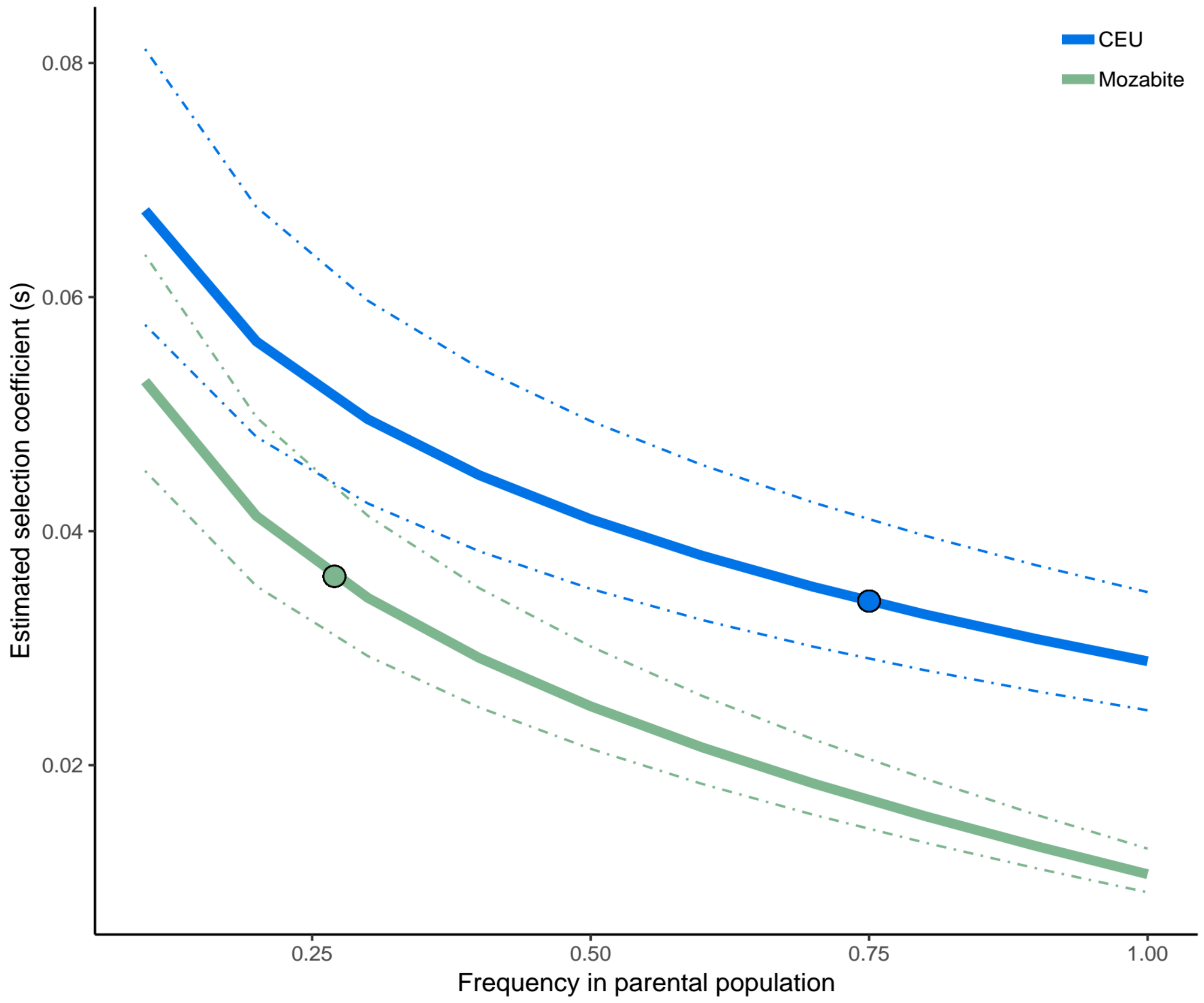


***Figure S17 - Estimated selection coefficient (s) of European LP T-13910 allele in the Fulani from Burkina Faso using the formula of Ohta and Kimura (1975).*** *CEU (in blue) and Mozabite (in green) were used as parental populations and the admixture fractions were based on the average ancestry proportions calculated by RFMix (Figure S4). For CEU as parental group, we fixed the admixture proportion to 13% and for the Mozabite to 32% (13% + 19%). The bold curves are calculated if we assume the admixture event to occur 63.0364 generations ago (Table S2), and dotted lines are if admixture event ± SD (10.695 generations ago) is considered. X-axis represents the T-13910 allele frequency in the parental population at the time of admixture. Dots are the selection coefficient calculations if the allele frequency of T-13910 in the parental group during the admixture event was the same as the allele frequency today.*
